# Integration of diverse bioactivity data into the Chemical Checker compound universe

**DOI:** 10.1101/2024.12.04.626832

**Authors:** Arnau Comajuncosa-Creus, Martino Bertoni, Miquel Duran-Frigola, Adrià Fernández-Torras, Oriol Guitart-Pla, Nils Kurzawa, Martina Locatelli, Yasmmin Martins, Elena Pareja-Lorente, Gema Rojas-Granado, Nicolas Soler, Eva Viesi, Patrick Aloy

## Abstract

Chemical signatures encode the physicochemical and structural properties of small molecules into numerical descriptors, forming the basis for chemical comparisons and search algorithms. The increasing availability of bioactivity data has improved compound representations to include biological effects, although bioactivity descriptors are often limited to a few well-documented molecules. To address this issue, we implemented a collection of deep neural networks able to leverage the experimentally determined bioactivity data associated to small molecules and infer the missing bioactivity signatures for any compound of interest. However, unlike static chemical descriptors, these bioactivity signatures dynamically evolve with new data and processing strategies. Here, we present a computational protocol to modify or generate novel bioactivity spaces and signatures, describing the main steps needed to leverage diverse bioactivity data with the current knowledge, as catalogued in the Chemical Checker (CC), using the predefined data curation pipeline. We illustrate the functioning of the protocol through four specific examples, including the incorporation of new compounds to an already existing bioactivity space, a change in the data pre-processing without altering the underlying experimental data, and the creation of two novel bioactivity spaces from scratch, which are completed in under 9 hours using GPU computing. Overall, this protocol offers a guideline for installing, testing and running the CC data integration approach on user-provided data, with the aim of extending the annotation presented for a limited number of small molecules to a larger chemical landscape.

**Key points:**

- The Chemical Checker is a large collection of processed, harmonized and integrated bioactivity signatures for over 1 million small molecules. Data are organized into 5 levels of increasing biological complexity and curation degrees: from chemical properties to clinical outcomes and from raw data representing explicit knowledge to embedded signatures inferred from observed bioactivity patterns.
- The Chemical Checker package provides a predefined and versatile framework to integrate user-provided data to the Chemical Checker universe of small molecules and generate novel customized bioactivity signatures.

**Key references:** Duran-Frigola *et al*. “Extending the small-molecule similarity principle to all levels of biology with the Chemical Checker.” Nature Biotechnology 38.9 (2020): 1087-1096.

Bertoni *et al*. “Bioactivity descriptors for uncharacterized chemical compounds.” Nature Communications 12.1 (2021): 3932.

## Introduction

Small molecules are excellent tools for probing biological functions and continue to be the cornerstone of pharmaceutical companies. Synthetic chemical compounds present a complex structural code that, for the most part, has not been explored by the principles of natural evolution. Many pharmacological compounds are imperfect human inventions with sub-optimal bioactivities that lack a clear connection to their chemical structure. Often, the only way to approach the biological characterization of a compound is to assume it will have comparable bioactivities to compounds with similar chemical properties. The so-called ‘similarity principle’ has been the driving force of drug discovery efforts^1^, and the measurement of compound similarities lays behind most of the approaches used to explore the vast drug-like chemical space (estimated in the range of 10^33^ molecules^2^). To assess such similarities, molecules are usually characterized using numerical fingerprints or descriptors (e.g.^3^), which encapsulate their main topological and physicochemical properties representing them in a format amenable for computational applications (e.g. QSAR)^4^.

In the last years, the extensive gathering and release of bioactivity data have shown that the similarity principle applies beyond standard chemical features, reaching to functional properties. For instance, small molecules that show similar sensitivity profiles across human tumor cell-lines or drugs that cause comparable side effects in patients tend to share mechanisms of action (MoA), even when they are structurally dissimilar^5–7^. Thus, bioactivity similarities offer an alternative means to functionally characterize small molecules, potentially providing insights closer to clinical observations and surpassing what is expected from merely inspecting chemical analogues^8^.

### The Chemical Checker

We recently developed the Chemical Checker (CC, **Fig 1a**), the largest collection to date of processed, harmonized and integrated bioactivity signatures for over 1 million compounds, whose biological effects had been experimentally determined^9^. Following the way in which we understand drug action, we organized data in 25 bioactivity spaces grouped in 5 levels of increasing biological complexity: from physicochemical and structural properties of small molecules (A) to the clinical effects they exert in human patients (E). In between, we collect the set of biological targets they bind (B), the perturbations they trigger in biological pathways (C) and the phenotypic effects they induce in cell-based assays (D). Unfortunately, bioactivity data become scarcer and more difficult to obtain as biological complexity increases (i.e. we have direct protein-binding data for ∼630k small molecules but only ∼6k compounds are annotated with clinical indications), meaning that bioactivity information remains limited for poorly characterized compounds. Indeed, out of the 10^33^ synthetically accessible drug-like molecules^2^, only a tiny fraction has experimentally measured chemical properties or bioactivities reported in public databases (i.e. ∼120M in PubChem and ∼2.4M in ChEMBL as of October 2024). To address this problem, we developed a collection of deep neural networks to infer CC bioactivity signatures for any compound of interest^10,11^, based on the observation that the different bioactivity spaces are not completely orthogonal, and thus similarities of a given bioactivity type (e.g. targets) can be transferred to other data kinds (e.g. therapeutic indications). Moreover, we showed that the inferred bioactivity signatures are useful to navigate the chemical space in a biologically relevant manner and improve the performance of biophysics and physiology activity prediction activities with respect to chemistry-only based classifiers^10^. In general, we believe that bioactivity signatures associated to small molecules have the power to reintroduce the biological complexity at the early stages of the drug discovery process, overcoming some of the problems associated with target-based approaches. For instance, using CC signatures, we were able to identify three compounds able to revert transcriptional signatures related to Alzheimer’s disease *in vitro* and *in vivo*^12^. Moreover, using CRISPR-Cas9 and shRNA perturbation experiments as templates, we also identified several small molecules that mimicked the phenotypic effects of biodrugs (e.g. daclizumab, ustekizumab, cetuximab, etc.), often through a different MoA that did not directly modulate the activity of their primary targets (e.g. IL-2R, IL-12 and EGFR)^9^, and targeted cancer proteins though to be undruggable^10^.

**Figure 1:**
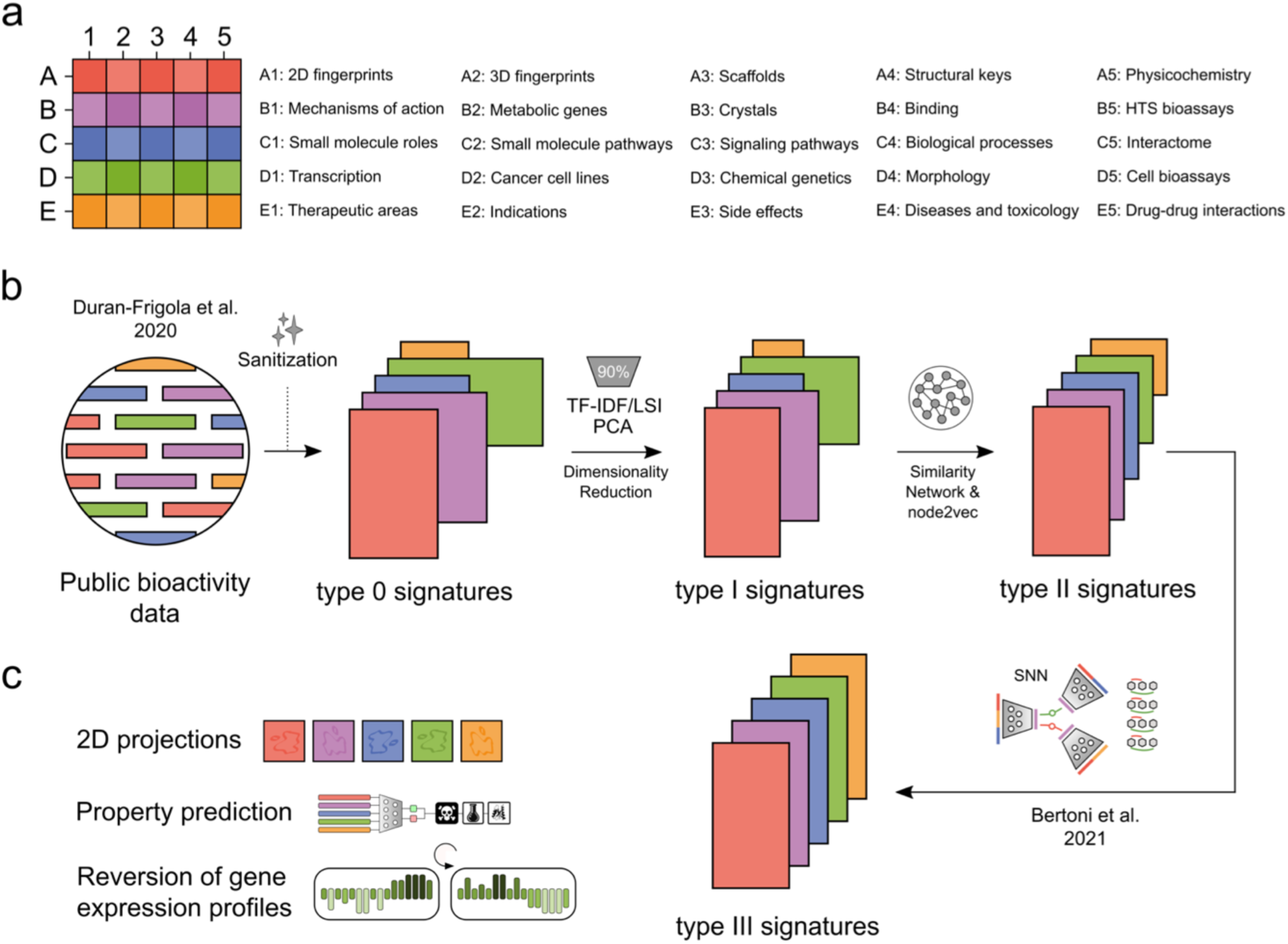
CC Overview. **a)** Organization of the 25 CC spaces. **b)** Computational pipeline to build the CC and procedure to integrate bioactivity data in the CC universe. In brief, bioactivity data is sanitized (type 0 signatures), compressed (type I signatures), harmonized (type II signatures) and integrated with all the other data within the CC (type III signatures). **c)** Compound signatures are used in a variety of tasks, such as the characterization and visualization of large small molecule libraries, the prediction of multiple properties (e.g. toxicity, physicochemical properties, protein binding, etc.) and the reversion of gene expression profiles associated with specific disease states, among many others.

The original 25 CC bioactivity spaces (**Fig 1a**) comprise data retrieved from publicly available databases at the time that were processed through a unified data curation and integration pipeline (see **Table S1** for a detailed list of the bioactivity databases integrated in the CC). Compound bioactivities are expressed in a vector-like format (i.e. signatures) and organized in increasing levels of abstraction: from raw data accounting for explicit knowledge (type 0 signatures) to inferred representations leveraging known bioactivity patterns (type III signatures). However, in the last years, new large-scale assays to capture biological responses to small molecule perturbations, and more efficient computational strategies to process them, are constantly appearing (e.g.^13–15^). Besides, there is an ever-growing number of laboratories and biotechnological companies that have their own in-house data and want to leverage it together with the bulk of known bioactivity information. Thus, in this manuscript, we present the complete computational protocol to generate novel bioactivity spaces and signatures, describing the main steps needed to leverage diverse bioactivity data with the Chemical Checker (type 0, type I, type II and type III signatures) using the predefined data curation and integration pipeline.

### Overview of the Chemical Checker data integration pipeline

The main input for the CC integration pipeline is a bioactivity data matrix encompassing compounds (rows, e.g. InChIKeys) and biological features (columns, unique strings: e.g. protein targets (B4), mechanisms of action (B1), clinical side effects (E3), etc.). Data might be categorical (e.g. binary), discrete or continuous.

#### Type 0 signatures

The very first step in the CC processing framework is to produce a sufficiently-curated and sanitized version of the raw bioactivity data (**Fig 1b**). To achieve this, features (columns) that are extremely sparse (i.e. occurring in few rows) or too prevalent (i.e. occurring in almost all rows) are removed from the matrix and those compounds (rows) showing no feature occurrences are discarded. In addition, not-available (NA) and infinite values are processed (median-imputed and max/min values, respectively) and low-Shannon entropy features are trimmed if the maximum number of allowed features is surpassed. Overall, type 0 signatures represent explicit knowledge, allowing direct interpretation while having variable dimensionality.

#### Type I signatures

The second processing step implies a mild compression of type 0 signatures typically retaining 90% of the original variance and thereby reducing the number of vector dimensions (**Fig 1b**). For discrete data, a term frequency–inverse document frequency (TF–IDF) transformation is used to weight features (columns) according to their information retrieval (i.e. more frequent features are less informative and thus become underweighted). After that, a Latent Semantic Indexing (LSI) dimensionality reduction technique is applied to the TF–IDF transformed data and LSI components explaining 90% of the original variance are kept to build type I signatures. For continuous data, the matrix is first scaled column-wise (median 0, median absolute deviation 1 and capped at ±10) and then processed through a PCA. Analogously to discrete data, PCA components explaining 90% of the original variance are kept to build type I signatures. Overall, type I signatures are better optimized for compound similarity calculations, although they no longer represent explicit knowledge and still have variable dimensionality.

#### Type II signatures

The main purpose of type II signatures is to retain similarities at type I signature level while harmonizing the length of vectors among CC spaces (i.e. converting signatures of variable dimensionalities to 128-length vectors, **Fig 1b**). First, a similarity network is built considering nearest neighbors (NN) at type I signature level for each compound (small molecules represent the nodes in the graph and their similarities are the edges). Then, the node2vec algorithm is run to create an embedding for each molecule (node), encoding intrinsic relationships within the similarity graph. In this way and, although not directly interpretable, type II signatures capture explicit and implicit similarities in the data. Additionally, the fixed length (128) makes the concatenation of descriptors across bioactivity spaces cost-effective, while ensuring that each bioactivity space contributes equally to the overall representation.

#### Type III signatures

The last step in the processing framework is the generation of type III signatures for the input compounds and for all the molecules in the CC, which capture and infer the similarity of the data at type I signature level (**Fig 1b**). The approach hinges on the insight that different CC spaces are not completely orthogonal and may show correlation to some extent. In short, for each CC space, a Siamese Neural Network (SNN) is trained over 10 million triplets of compounds (anchor, positive and negative molecules sampled at type I signature level) representing these input molecules by concatenating all available type II signatures. After being validated, the SNN generates novel signatures (type III) that retain the observed similarities at type I signature level and accounts for compound data present in every other bioactivity space of the CC, allowing the inference of a bioactivity signature even in the absence of data for the molecule of interest. It is worth mentioning that the generation of type III signatures is usually the most computationally expensive step in the CC data integration pipeline, and strongly benefits from parallelization and GPU computing (see *Materials: Hardware*).

Since most properties and similarities at type III signature level are inferred, each of them comes together with a measure of confidence called ‘applicability’ (from 0 to 1). Applicability scores evaluate the distance to training-set signatures, the robustness of the predicted signatures and the expectancy a priori given the known data. For further details about signature inference and confidence score, we refer the reader to the original publication^10^.

#### Additional remarks

The outlined pipeline includes numerous distinct steps, each possessing its own set of default options and parameters. While the user can modify the vast majority of parameters, the whole point of the computational pipeline –and where we put most emphasis– is in the full automation. In fact, the approach works end-to-end, and it is very robust to very different data shapes and types (sparse, long, wide, dense, etc.). Thus, we recommend to use the by default parameters, but give the possibility to the user to adapt them if new uses of the CC signatures might arise. For additional details on such parameters and further clarification of the mentioned steps we refer to the original publications^9,10^ and our regularly maintained GitLab repository (https://gitlabsbnb.irbbarcelona.org/packages).

#### Interpreting the results: diagnosis plots

To assess and monitor data flow at each step of the CC integration pipeline, we have designed several analyses to evaluate the results in a systematic manner. In brief, for each data matrix (compound signatures), a set of diagnosis plots is generated to illustrate the main characteristics of the data. This section includes a comprehensive explanation for each plot.

Examples for diagnosis plots are found in **Fig S1, 4, 7 and 10**. From left to right and from top to bottom:

- Image: heatmap representing the data matrix. Rows are compounds (keys, max. number set to 100), columns are features and colors represent actual values from minimum (blue) to maximum (red).
- Values distribution: density distribution (y-axis) of data values (x-axis) from minimum (blue) to maximum (red).
- 2D projection: tSNE (t-distributed Stochastic Neighbor Embedding) representation of the data, colored by density. The maximum number of subsampled compounds included in the projection is 10 thousand.
- Euclidean distribution: density distribution (y-axis) of pairwise Euclidean distances (x-axis). The maximum number of selected compound pairs is set to 10 thousand.
- Values by feature: numerical values (y-axis) of each feature (x-axis).
- Values by key: numerical values (y-axis) of each compound (key, x-axis).
- Redundancy: for each signature in a non-redundant set of signatures (x-axis), number of compounds (keys, y-axis) having the same exact signature.
- Cosine distribution: density distribution (y-axis) of pairwise cosine distances (x-axis). The maximum number of selected compound pairs is set to 10 thousand.

The full version of the diagnosis plots provided by chemical checker comprise additional analysis and comparisons to other CC spaces. Examples of the full version are found in Fig S1, 5, 8 and 11. The new plots in the full version are described below from left to right, top to bottom:

- Confidence: Density plot of normalized confidence values for the signatures. Mean values above 0.6 indicate reliable signatures.
- 2D projection colored by confidence value: t-SNE representation of the data, colored by their assigned confidence values.
- CC ranks agreement: Distribution of the agreement between compounds neighbors rankings. The lower the score, the lower the agreement among the compounds neighbors rankings
- 2D projection colored by CC ranks agreement: t-SNE representation of the data, colored by their neighbor rank agreement.
- Outliers: It represents the anomaly score for each signature Signatures with scores above 0 are considered outliers.
- Key Coverage: Proportion of compounds shared across different CC space datasets. Any given compound may be present in all or some CC spaces (1-25, x-axis).
- 2D projection colored by key coverage: t-SNE representation of the data, colored by the compound coverage. Blue indicates higher compound presence across CC spaces.
- Cluster sizes: Proportion of keys assigned to each cluster. We obtain the clusters by computing the distances between the signatures and then apply DBScan method to find the neighborhoods. Top 20 clusters are highlighted in the plot
- Top clusters:2D representation of the top 20 clusters.
- MoA: Recapitulation of the nearest neighbors in the B1 space (MoA) using the current signature. This recapitulation is quantified by AUROC (Area Under the Receiver Operating Characteristic) values, indicated in the figure title. Values around 0.5 indicate random results
- ATC: Recapitulation of the nearest neighbors in the E1 space (therapeutic areas; ATC) using the current signature.
- Keys / Features dimension: Number of compounds and features across all CC spaces for the current signature. The values were normalized using log10.The current CC space dot is highlighted with black border and white background.
- CC wrt Sign: Number of keys of the current space that overlaps with the keys of all the other spaces colored according to the CC levels color legend. Numbers (ranging from 1 to 5) correspond to each space per level.
- Sign wrt CC: Overlap among the keys of all the other spaces and the keys in the target space.
- ROC across CC: Recapitulation of the nearest neighbors of all CC spaces. area under ROC curve values summarize the performance. The AUC values are sorted in descending order (higher values better recapitulation) and colored according to the CC levels color legend. Numbers (ranging from 1 to 5) correspond to each space per level.

Generating diagnosis plots is a straightforward task with the CC package (please see *Procedure: Step 10*). Further information on diagnosis plots and additional data analysis (e.g. recapitulation of MoA and ATC) are found in our Gitlab. More importantly, **Table S2** provides specific guidance on how to interpret the results for each panel of the diagnosis plots.

### Applications of the method

To comprehensively illustrate the use of the protocol to build bioactivity spaces and signatures associated to small molecules, we adapt it to four distinct scenarios: i) introducing additional data into an existing CC bioactivity space, ii) creating a new version of an existing CC bioactivity space with a different strategy of input data processing, iii) creating a CC bioactivity space from a novel data type that fits the actual CC universe (i.e. human-derived data) and iv) creating a CC bioactivity space from a novel data type that does not fit the actual CC universe (i.e. measuring compound effects on bacterial species). All 4 scenarios are briefly described in *Experimental design* and fully presented in the *Anticipated results* section. Overall, in all cases, novel bioactivity data is processed, sanitized, compressed, harmonized and integrated in the CC universe of small molecule signatures. Moreover, bioactivity signatures enable the exploration and visualization of large chemical libraries in a biologically relevant manner as well as the prediction of compound properties such as toxicity, protein binding and the ability to revert gene expression profiles associated with certain disease states, among others (**Fig 1c**).

### Comparison with other methods

Chemical descriptors are at the core of cheminformatics. Indeed, there are many strategies aimed at encapsulating the structure of drug-like compounds in a format amenable for computational applications. Extended connectivity fingerprints (ECFPs) represent the most popular approach to do so, due to their high interpretability, low computational cost and strong performance in various prediction tasks^16^. Other classical and popular representations include substructure-based fingerprints^17^, Murcko scaffolds^18^ and, more recently, SMILES- and graph-based embeddings built upon transformer architectures (e.g.^19,20^). However, as stated in the *Introduction*, bioactivity descriptors offer an alternative (and arguably more clinically relevant) perspective to navigate the chemical space in the search for new bioactive compounds with desired pharmacological properties. To our knowledge, there is no alternative strategy for processing and integrating user-provided biological data in response to chemical perturbations to generate customized bioactivity descriptors.

### Experimental design

In this manuscript, we present and run the described computational protocol for four distinct tasks, organized in increasing level of conceptual and technical complexity.

1. Extend a preexisting CC space with additional bioactivity data: **B1.002** space.
2. Rebuild a preexisting CC space with a different strategy of processing the raw data: **D1.002** space.
3. Create a new CC space from a novel data type that fits the current CC universe (i.e. human-derived data): **D6.001** space.
4. Create a new CC space from a novel data type that does not fit the current CC universe (i.e. measuring compound effects on bacterial species): **M1.001** space.

The CC nomenclature to define the different bioactivity spaces first notes the space (i.e. A1-n, B1-n, etc) and then numbers the dataset or processing protocols used (i.e. 001, 002, etc).

### Expertise needed to implement the protocol

The protocol can be implemented by anyone having basic experience with the Bash Command Line (mainly for the local installation of the CC) and basic scripting knowledge of the Python coding language (to actually run the designed protocol). Additionally, familiarity with high-performance computing (HPC) cluster and queuing systems is recommended in case of i) handling of large amounts of data or ii) running the last integration step of the pipeline, for which parallelization or GPU computing is usually helpful (see *Procedure: Step 9* and *Limitations*).

### Limitations

The CC data integration pipeline has some drawbacks and limitations that need to be acknowledged before use. First, as all frameworks designed to process external user-provided data, the integrity of the final outcome is ultimately tied to the quality of the input data: suboptimal, noisy or otherwise compromised data can significantly degrade the results and produce unreliable signatures. To help the user evaluate their signatures, the diagnosis plot provides a quantification of biological and chemical signals recapitulated by the newly generated spaces, intended to provide insights on the biomedical information captured by the user data.

It is also important to consider that input datasets having a small overlap with the CC compound universe may result in type III signatures mostly based on the chemical properties of the small molecules, since the lack of additional bioactivity data might impact SNN training. In line with this, it is likely that small molecules with no reported bioactivity in a specific CC space are eventually characterized by the so-called *null signatures*, corresponding to the average signature derived from the training set. To partially control this effect and handle additional problematic issues (see *Introduction: type III signatures*), every inferred CC type III signature is accompanied by an applicability score that measures the overall degree of confidence in the compound signature. In this way, such applicability scores make it possible to filter out compounds that have unreliable signatures, potentially leading to more accurate and trustworthy results.

Another limitation of most CC bioactivity signatures, similar to many machine and deep learning applications^21,22^, is the lack of direct interpretability. Only type 0 signatures are directly explainable, since signature elements (columns) correspond to well-defined biological features (e.g. chemical substructures, binding targets, clinical side effects, etc.). However, type I, II and III signatures are abstract molecular representations for which experimental data (biological features) cannot be deconvoluted. In any case, we have already shown that CC signatures are useful across various tasks, and it is indeed possible to achieve a nuanced degree of interpretability by identifying which bioactivity signatures are more valuable and relevant for the predicted outcomes^10^.

Finally, another potential limitation of the presented pipeline is the relatively high computational cost associated with the generation of type III signatures. This step integrates all available bioactivity signatures in the CC and benefits from memory availability and extensive parallelization and GPU computing. We have designed a twin version of the type III signature fit method to allow the execution of this step in a predefined HPC cluster (see *Procedure: Step 9*). It is worth mentioning that computational cost is usually not a problem in the other steps of the protocol, as illustrated in the *Anticipated results* section for various distinct input datasets. In addition, sufficient free hard disk space is essential for successfully downloading CC data and running the computational protocol smoothly. Overall, we encourage users to free +150GB of their machines (see *Troubleshooting*).

## Materials

### Hardware

Workstation with 4 CPUs and 16GB of RAM. Optionally, a computational cluster can be used to speed up the ‘signaturization’ process (mainly of type III signatures), exploiting wider task parallelization and GPU computing as well as higher memory availability in case of inputting a large dataset to the CC integration pipeline.

### Software

Linux or Linux-emulating operative systems. The CC package needs to be installed following the steps detailed in *Procedure: Step 1*. A Singularity image is built during the installation process, including all required software packages and their dependencies to run the CC pipeline in a safe environment. For other systems, the creation of the Singularity image might not be supported, and it might require a virtual environment. In addition, Apple computers with M processors (e.g. M1, M2 or M3) may face unanticipated issues.

### Other

In order to generate type III signatures, the user needs to get all type II signatures as well as additional dependencies from the current CC version. Signatures can be downloaded as plain data files directly from https://chemicalchecker.com/downloads/root, using the python code we provide to do so (check *Procedure step 3*), or accessing through the CC REST API (https://chemicalchecker.com/help, bottom of the page).

## Procedure

### Preliminary steps

#### 1. Setting up the CC environment ∼30 min

[**CRITICAL STEP**] The installation of the CC in a local computer is carried out following the instructions provided in:

https://gitlabsbnb.irbbarcelona.org/packages/chemical_checker

In short, we need to i) install Singularity (version>3.6), ii) clone the CC repository to a local directory and iii) run the setup script (*sudo* permissions are needed). After that, we need to iv) adapt the CC configuration file to fit the user’s CC environment and, finally, v) clone the CC protocols repository to implement the work described in this article (see *Anticipated results*).

i. Installing Singularity [**TROUBLESHOOTING**].

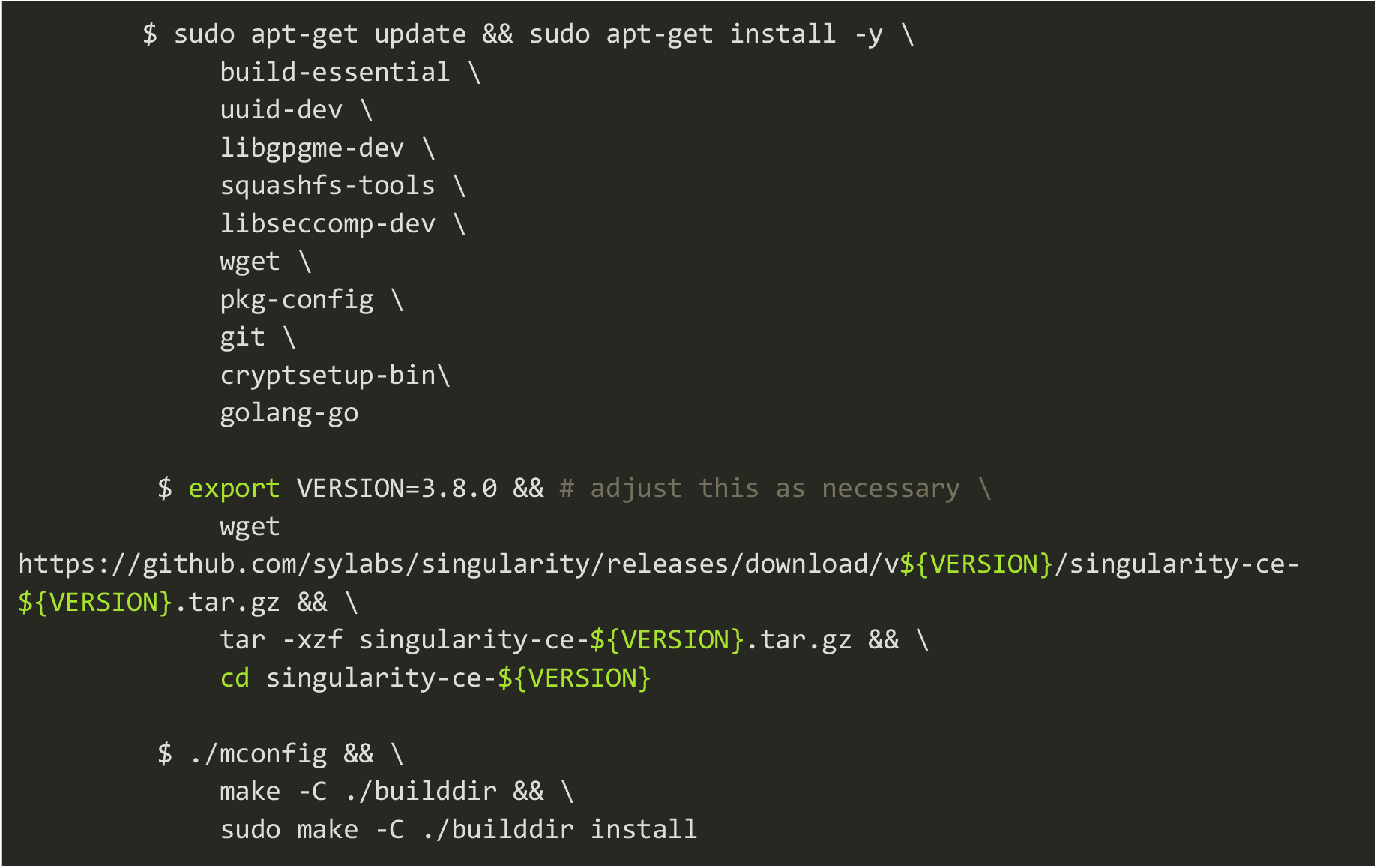
ii. Cloning the CC repository to a local directory named *PATH_TO_CC* [**TROUBLESHOOTING**].

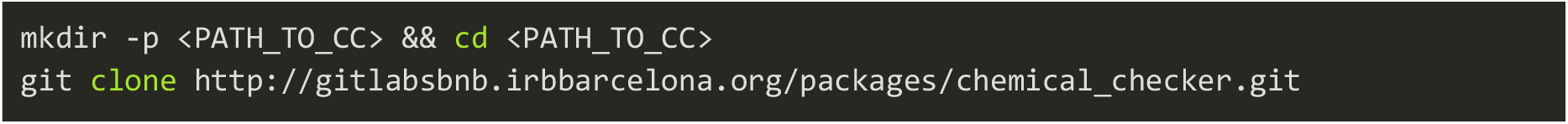
iii. Running the CC setup script [**TROUBLESHOOTING**].

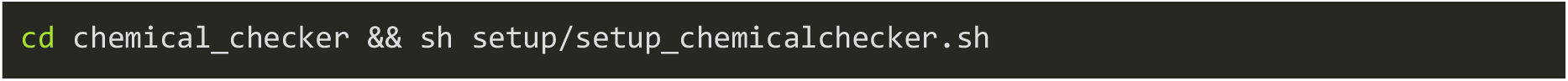
iv. Manually editing the CC configuration file located at *PATH_TO_CC/chemical_checker/setup/cc_config.json*, referred to as *PATH_TO_CC_CONFIG*. In this file, the user must specify the variables *CC_REPO*, *CC_TMP* and *SINGULARITY_IMAGE*.

- *SINGULARITY_IMAGE* should point to the CC singularity image created when running the CC setup script (see Procedure step 1.iii), usually located in *$HOME/chemical_checker/cc.simg*, *$HOME* being the user’s home directory.
- *CC_REPO* should point to the chemical_checker directory inside *PATH_TO_CC (i.e., PATH_TO_CC/chemical_checker*, see Procedure steps 1.ii and 1.iii).
- *CC_TMP* should point to a local directory, defined and created by the user, where CC outcome files will be stored. This directory is referred to as *PATH_TO_CC_TMP*.
v. Cloning the CC protocols repository to a local directory referred to as *PATH_TO_CC_PROTOCOLS*. This repository includes all the notebooks necessary to implement the analyses presented in this manuscript.

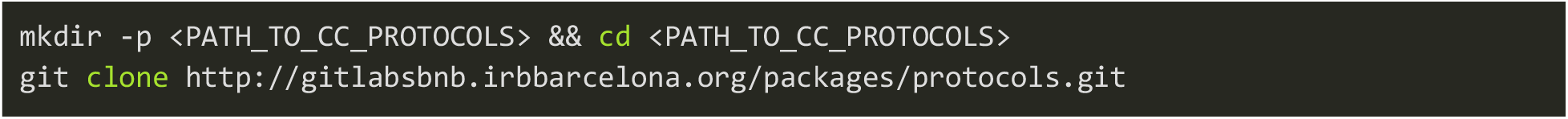

In brief, the repository contains four Python notebooks corresponding to the four tasks implemented herein (see *Anticipated results*), along with the raw bioactivity data required to build the new CC spaces. The following points in this section (i.e. Procedure) provide an overview of the individual steps needed to generate new CC spaces and signatures.

**[TROUBLESHOOTING]**

#### 2. Running a Jupyter Notebook with the CC image <1 min

We run a Jupyter Notebook in the CC protocols directory within the CC image by executing the following command (which is an alias to run the *setup/run_chemicalchecker.sh* file from the cloned CC repository) [**TROUBLESHOOTING**]:

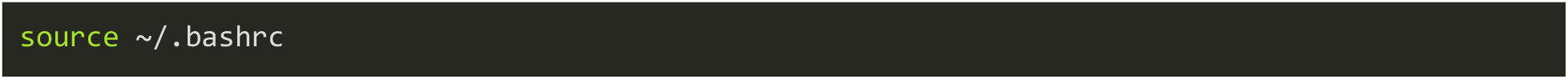

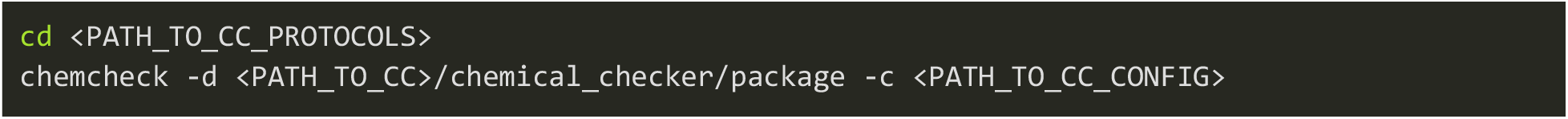

All notebooks needed to build the four spaces described in this manuscript are found in *PATH_TO_CC_PROTOCOLS*/*protocols*/*notebooks*.

[**CRITICAL STEP**] Once the Jupyter Notebook is up and running, it is essential to import the CC package itself as well as all necessary external libraries to run the analyses (e.g. numpy, see examples in *Anticipated results* and *Supplementary Information*). Before that, we also need to point the environment variable *CC_CONFIG* to the local CC configuration file (see *Procedure step 1.iv*). In addition and, after importing the CC package, we can specify the level of log outputs printed along the procedure. This piece of code is already included in all the exemplary notebooks.

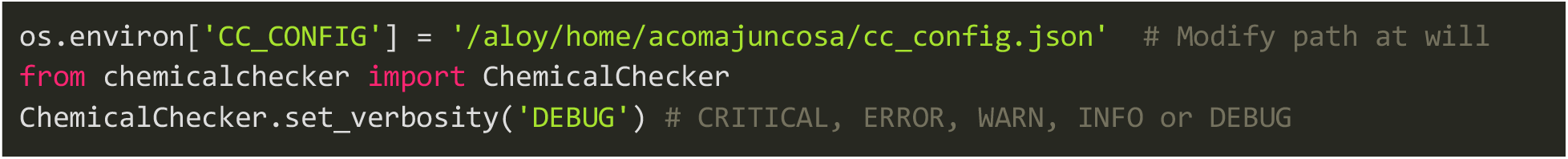

#### 3. Gathering CC data ∼45 min

To successfully implement all CC functionalities illustrated in this work, the user needs to gather the CC bioactivity data from our web servers (e.g. type II signatures, see Procedure 10). To illustrate the downloading process, we provide exemplary code that programmatically downloads the CC signatures, although the user can download individual files containing bioactivity signatures manually (https://chemicalchecker.org/downloads/signatureX, X being 0, 1, 2 or 3). In brief, data are first downloaded in a compressed tar.gz format and then uncompressed in the directory PATH_TO_DATA, as shown in the notebook Download_Data.ipynb included in the CC protocols repository (see code below). Since the CC is subject to yearly-carried updates, the size of the downloaded files may increase, as does the amount of publicly released bioactivity data. As of October 2024, the overall process takes 15 minutes to download and 30 minutes to decompress, requiring around 65 GB of free disk space and a stable wired internet connection (e.g Ethernet). For further details on what specific CC data are needed to implement each CC functionality, please see the *Supplementary Information*.

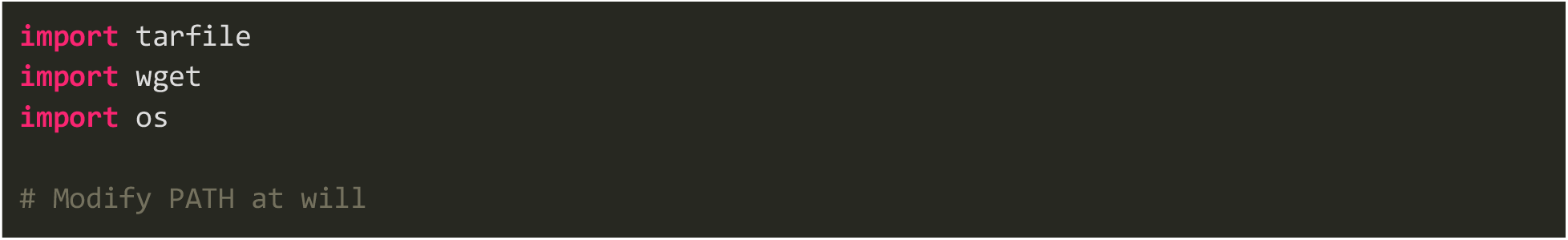

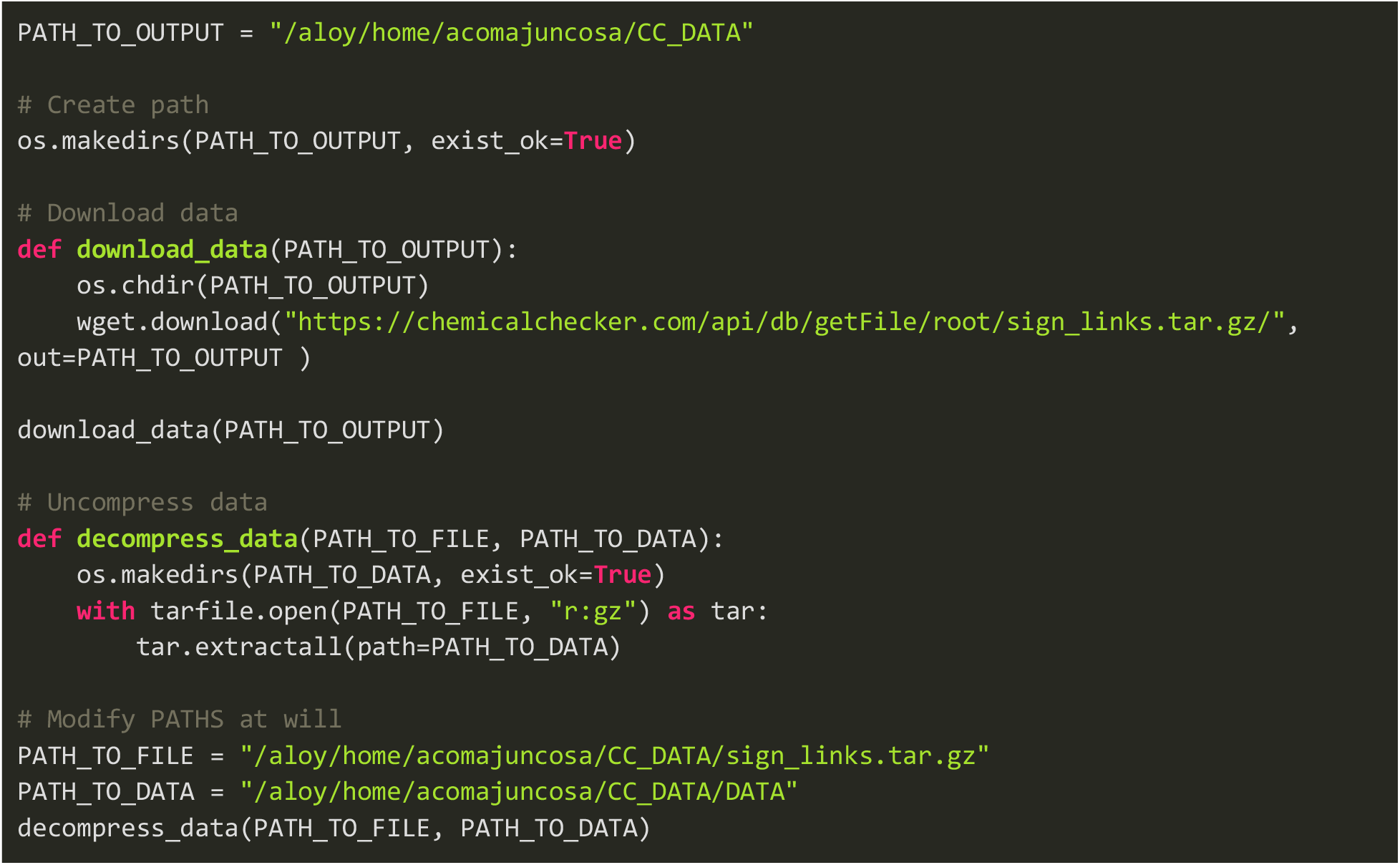

### Main steps

While preliminary steps are meant to be executed only once, the main steps of the protocol need to be run for each newly created CC space. The specific code developed to perform each individual task (see *Experimental Design*) is included in the CC protocols repository (see *Procedure step 1.v*). As an example, the code shown in this section corresponds to the new M1.001 space (i.e. task 4). The overall timing of the main steps fully depends on the type and size of input data. For the sake of completeness, we specify the timing of all main steps for the 4 distinct tasks in the *Timing* section.

#### 4. Creating a CC instance

A new CC instance is created with the directory chosen to allocate the results (*local_cc_dir*). Additionally, we also need to specify the location of existing CC data (*PATH_TO_DATA* see *Procedure: Step 2*). [**TROUBLESHOOTING**]

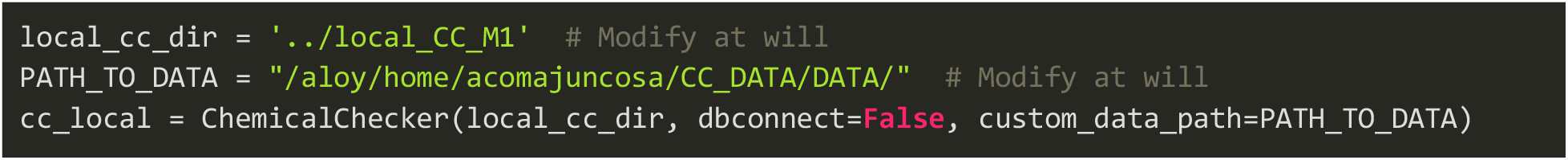

The typical CC folder architecture will be created inside *local_cc_dir*. The full directory will contain the computed signatures (from 0 to III) for the corresponding small molecules, while the reference set will include a non-redundant subset of the data, computed using the distance matrix among all compounds.

#### 5. Loading raw data [**CRITICAL STEP**]

To start the signaturization process, we first need to load a preprocessed version of the raw bioactivity data. Depending on the format of the data, the loading process may differ significantly. In Anticipated results and Supplementary code, we have included examples in which the data is located in a CSV file and in a H5 file (see the code below for loading the input csv data file for the new M1.001 space). Essentially, the starting data must consist in a well-defined bioactivity matrix (e.g. a pandas dataframe) containing N compounds (rows, indices) and M biological features (columns). Note that the code needs to be adapted in case of using user-defined data. For all the examples presented in this manuscript, we also provide the code to download the data from our online repository and store it in the corresponding data directory (not shown here) to be then loaded (shown below).

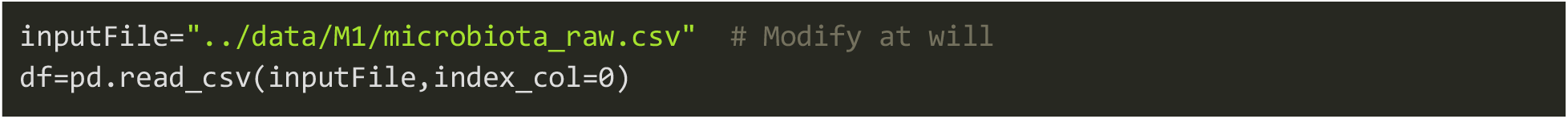

#### 6. Generating type 0 signatures

Once the bioactivity data is loaded, we can generate type 0 signatures with the following code block. It is important to mention that we first need to define a *dataset* name in a particular format specifying the CC space the data will be dumped in (e.g. M1.001, please see *The Chemical Checker* section). For further information about the generation of type 0 signatures, please see *Overview of the Chemical Checker data integration pipeline: Type 0 signatures*). [**TROUBLESHOOTING**]

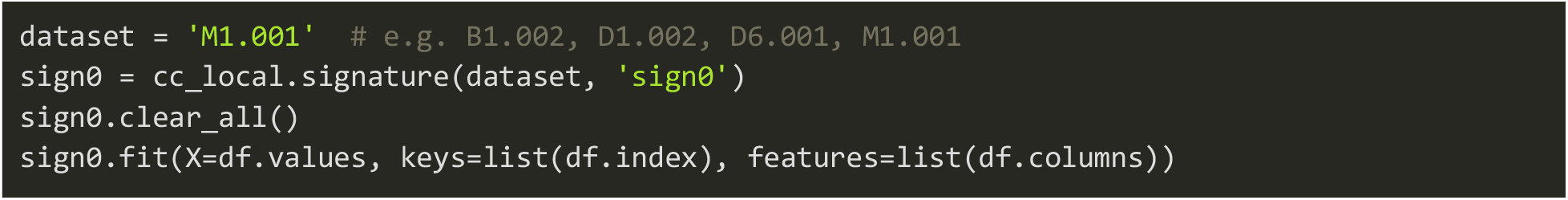

#### 7. Generating type I signatures

After obtaining type 0 signatures, we proceed to the generation of type I signatures. The code is straightforward and analogous to the previous step. Additionally, we also need to precompute compound nearest neighbors at type I signature level. For further information about the generation of type I signatures and the calculation of neighbors, please see *Overview of the Chemical Checker data integration pipeline: Type I signatures and Type II signatures*). [**TROUBLESHOOTING**]

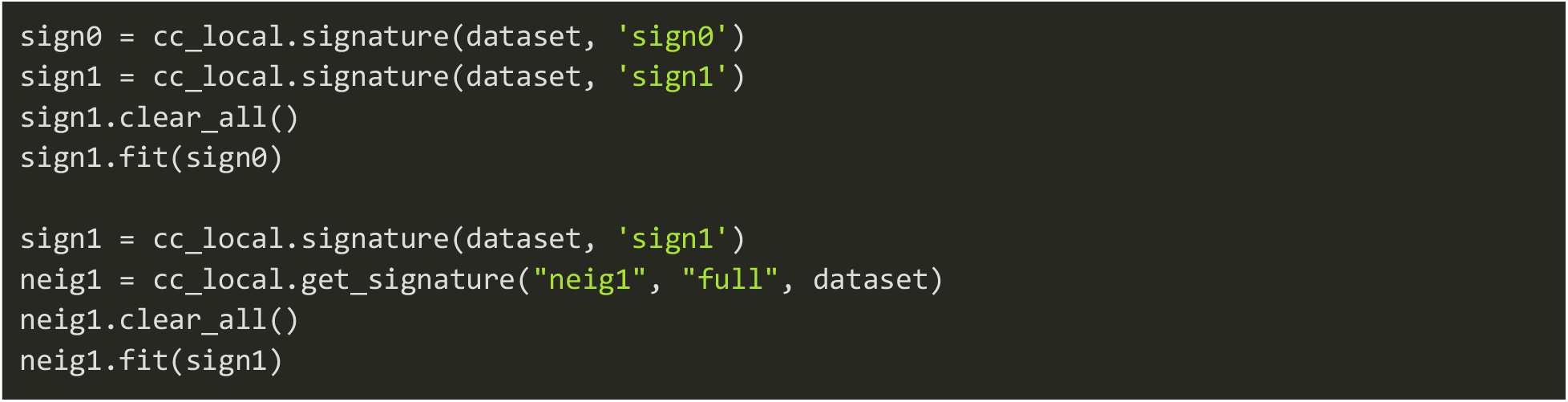

#### 8. Generating type II signatures

Once we have type I signatures and neighbors calculated for our compound set, we can proceed to the generation of type II signatures. As seen in previous steps, the code is straightforward and follows the same architecture as before. For further information about the generation of type II signatures, please see *Overview of the Chemical Checker data integration pipeline: Type II signatures*). [**TROUBLESHOOTING**]

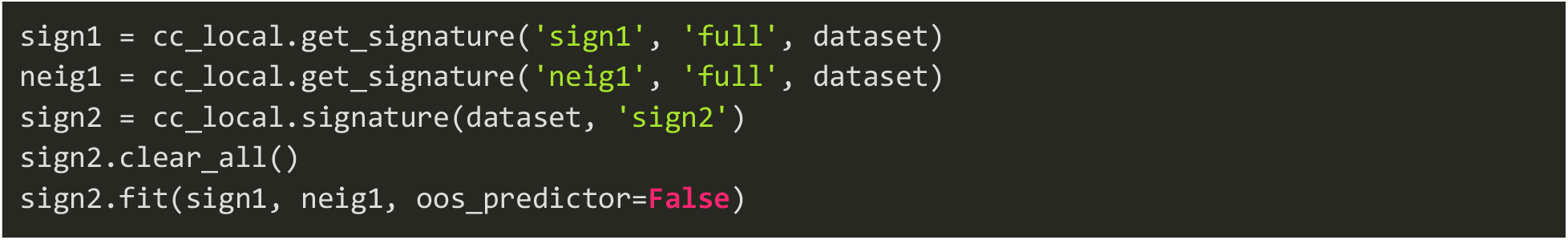

#### 9. Generating type III signatures

The last step in the signaturization process is the calculation of type III signatures. We first collect all type II signatures from the existing CC spaces along with type I and type II signatures for the extended or newly created space. Since the calculation of type III signatures is computationally expensive (see *Limitations*), we run it on an HPC cluster to enable CPU parallelization or GPU computing. Indeed, we encourage users to run the generation of type III signatures in their own HPC cluster. Otherwise, it can be run locally in the same manner that previous steps, although the overall calculation may take longer to complete. Additionally, when creating a new CC space, it is important to check the overlap between the input dataset and the CC universe of small molecules as poor overlaps between both sets may eventually result in unreliable type III signatures (see *Limitations*). Furthermore, chemical signatures (A1, A3, A4 and A5) are automatically calculated for all those compounds that are not included within the CC universe (*complete_universe=’fast’*). If the user also wants to include A2 signatures, a particularly slow process due to the generation of 3D conformers, the *complete_universe* parameter needs to be set to ‘*full’*. Finally, we encourage the user to provide the corresponding mappings between the input InChIKeys and the molecule InChIs in a Python dictionary. Otherwise, the pipeline will query online repositories (e.g. PubChem) to derive the mappings, which may significantly slow down the entire process. The code below, as all the code shown in the Procedure section, corresponds to the case example from *Anticipated results: Task 4* (M1.001) and it needs to be adapted to every particular exercise. For additional examples, please see *Supplementary code: Tasks 1-3*. [**TROUBLESHOOTING**]

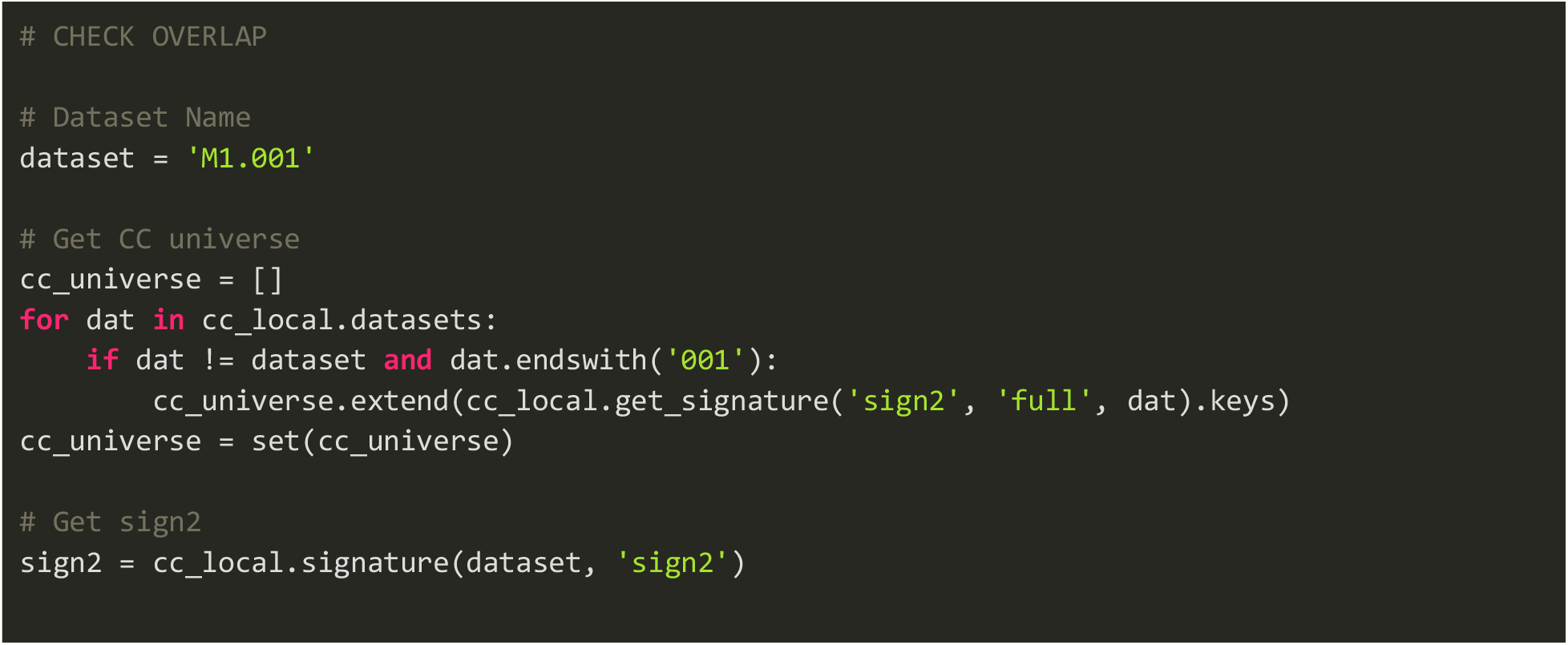

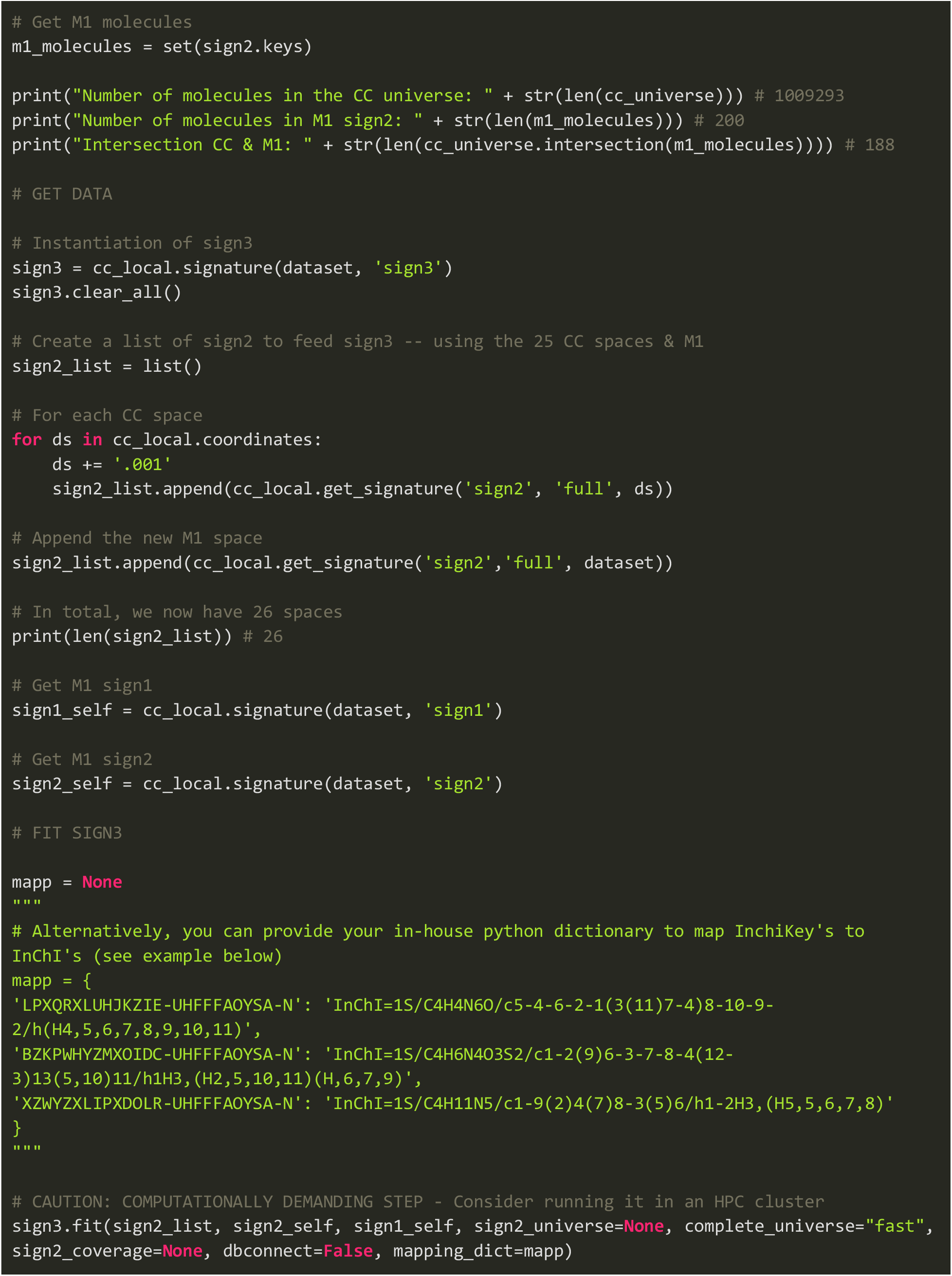

#### 10. Generating diagnosis plots

Finally, to visualize the flow of the data along the integration pipeline, we can generate diagnosis plots for all the newly obtained signatures (0, 1, 2 and 3) with the code provided below (replacing *X* by 0, 1, 2 or 3, depending on the signature type under analysis). For further information about the generation and interpretation of diagnosis plots, please see *Overview of the Chemical Checker data integration pipeline - Interpreting the results: diagnosis plots*). [**TROUBLESHOOTING**]

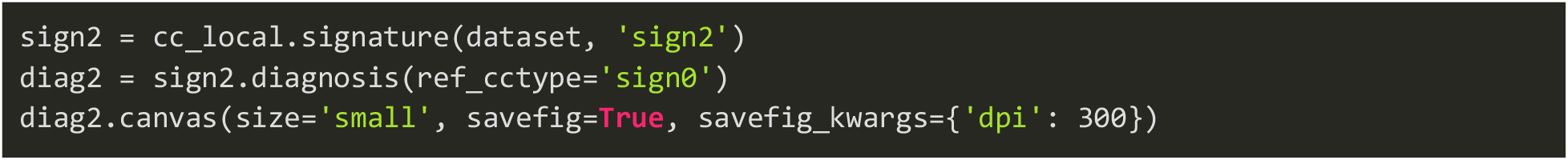

## Anticipated results

In this section, we showcase the execution of the CC data integration pipeline in four different scenarios, as briefly described in the *Experimental design* section (**Fig 2**). For each task, we first introduce the nature of the raw bioactivity data (*Dataset description*) and we then present and discuss the results obtained along the pipeline (*Signaturization results*). The implemented procedure is detailed in *Procedure: Main steps*, and the individual code adapted to each task is included in *Supplementary Code* as well as in the example notebooks located in our Gitlab.

**Figure 2:**
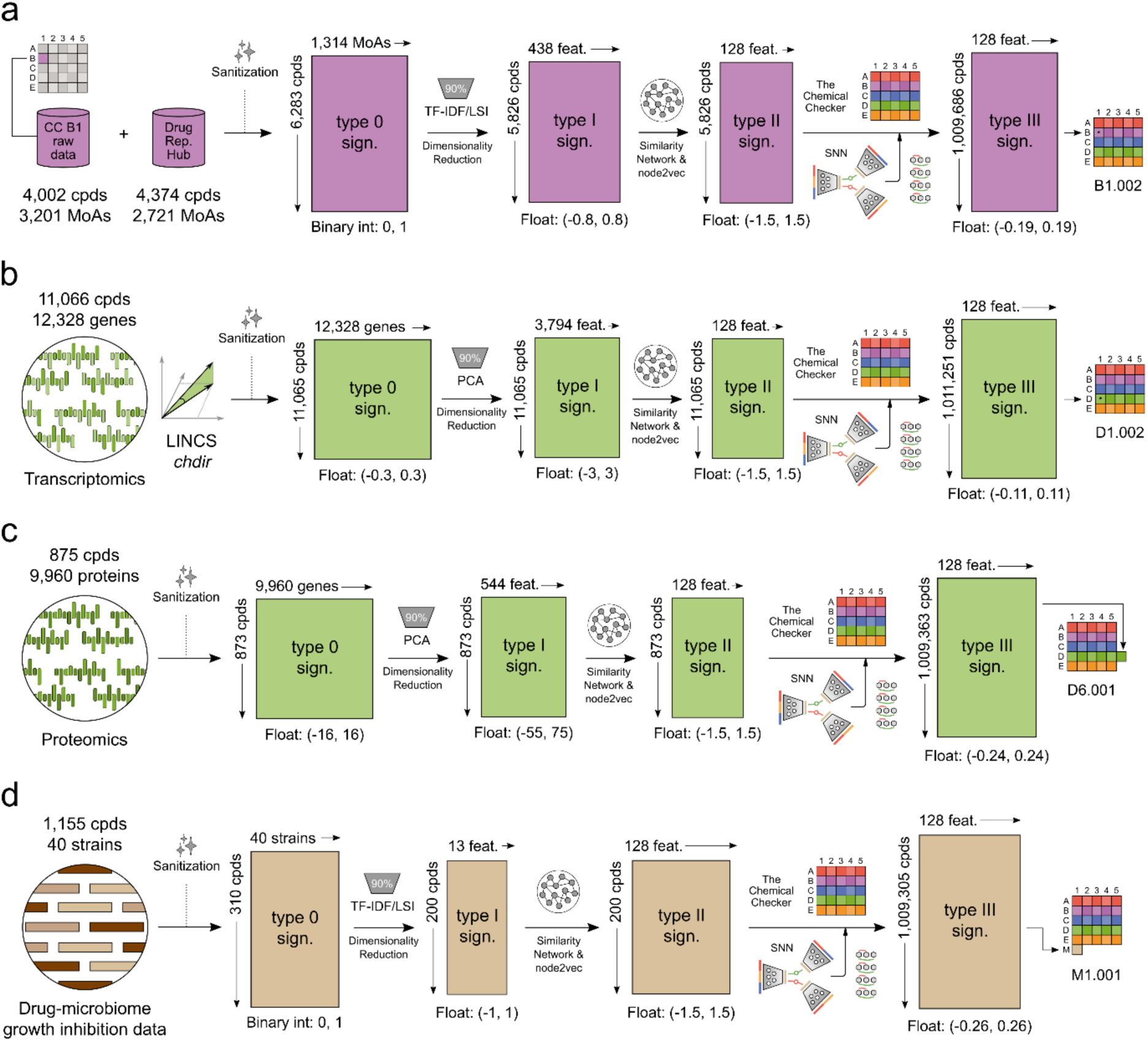
CC Protocols case examples. All steps along the pipeline are detailed in *Procedure* and the results are explained and discussed in the *Anticipated Results* section. **a)** Extending a preexisting CC space with additional bioactivity data (*Anticipated Results: Task 1*), **b)** Rebuilding a preexisting CC space with a different strategy of processing the raw data (Anticipated Results: Task 2), **c)** Create a new CC space from a novel data type that fits the current CC universe (i.e. human-derived data) (Anticipated Results: Task 3) and **d)** Create a new CC space from a novel data type that does not fit the current CC universe (i.e. measuring compound effects on bacterial species) (Anticipated Results: Task 4).

### Task 1: Extending a preexisting CC space with additional bioactivity data

#### Dataset description

The Drug Repurposing Hub^23^ is a centralized, comprehensive and curated collection of FDA-approved drugs, preclinical tool molecules, and clinically failed compounds for which safety has already been proven. All compounds are annotated with their chemical structures together with information about their mechanisms of actions (MoAs), approved disease indications and clinical trial status, if available.

The original CC B1 space (B1.001, MoA) was built using available data from ChEMBL (v.22) and DrugBank (v.4). The main purpose of this exercise is to extend the B1 space (namely B1.002) with the data from the Drug Repurposing Hub, accounting for 4,374 compounds and 2,721 mechanisms of action.

#### Signaturization results ∼ 8 hours

First, we observed that the overlap between the original (ChEMBL and DrugBank) molecule-MoA pairs and those coming from the Drug Repurposing Hub was low, with only 3,342 out of the 11,971 new pairs already annotated in the original CC B1.001 space, which shows the value of incorporating this dataset. We merged all data (6,944 compounds and 4,626 mechanisms of action) and started the signaturization process. 3,312 seldom occurring MoAs and 661 compounds exhibiting null bioactivity profiles were filtered out in the first processing step (*min_feature_abs* and *min_keys_abs* parameters, respectively). Eventually, we obtained type 0 signatures for 6,283 compounds with 1,314 mechanisms of action. As illustrated in the diagnosis plots, the new B1.002 type 0 signatures were binary and highly redundant (**Fig S1a**). In the next step, we reduced data dimensionality and obtained type I signatures for 5,826 compounds, with 438 PCA components capturing 90% of the variance. We then harmonized the number of features to match the CC data and generated type II signatures for these 5,826 molecules (128 features). Finally, we integrated the new B1.002 space together with the rest of the CC spaces and generated type III signatures for all 1,009,686 compounds (5,826 and 1,009,293 in B1.002 and the CC universe, respectively, with an overlap of 5,433 compounds).

An important consideration is that similarities at the raw data level (type 0 signatures) may be partially diluted along the data contextualization pipeline (e.g. at type III signature level), when the data is eventually compressed and integrated with all the bioactivity data already contained in the CC. To further illustrate which spaces may be more affected in this regard, we have designed an evaluation process in which, for each new CC space and signature type, we assess the ability to recapitulate NN from a signature type at a lower level (**Fig 3a**). First, for the newly created B1.002 space, we defined a cutoff cosine distance for all signature types (0, 1, 2 and 3, i.e. 0, I, II and III) at a pvalue of 0.01 (10,000 randomly selected compounds x 10 subsamples, considering the average value). Then, for each combination of signA-signB, we evaluated the ability of signB to recapitulate NN defined at signA level with a pvalue of 0.01 (for each combination, 2,500 random molecules, 5 subsamples; totaling 6 combinations of signature type pairs). In the case of the B1.002 space, we observed that type III signatures were able to greatly recapitulate similarities observed at the raw bioactivity data level (type 0 signatures, AUROCs∼0.85, **Fig 3b**). In addition, we also observed that applicability values associated with type III signatures were fairly in line with the values obtained for the original CC bioactivity spaces (**Fig 3c**).

**Figure 3:**
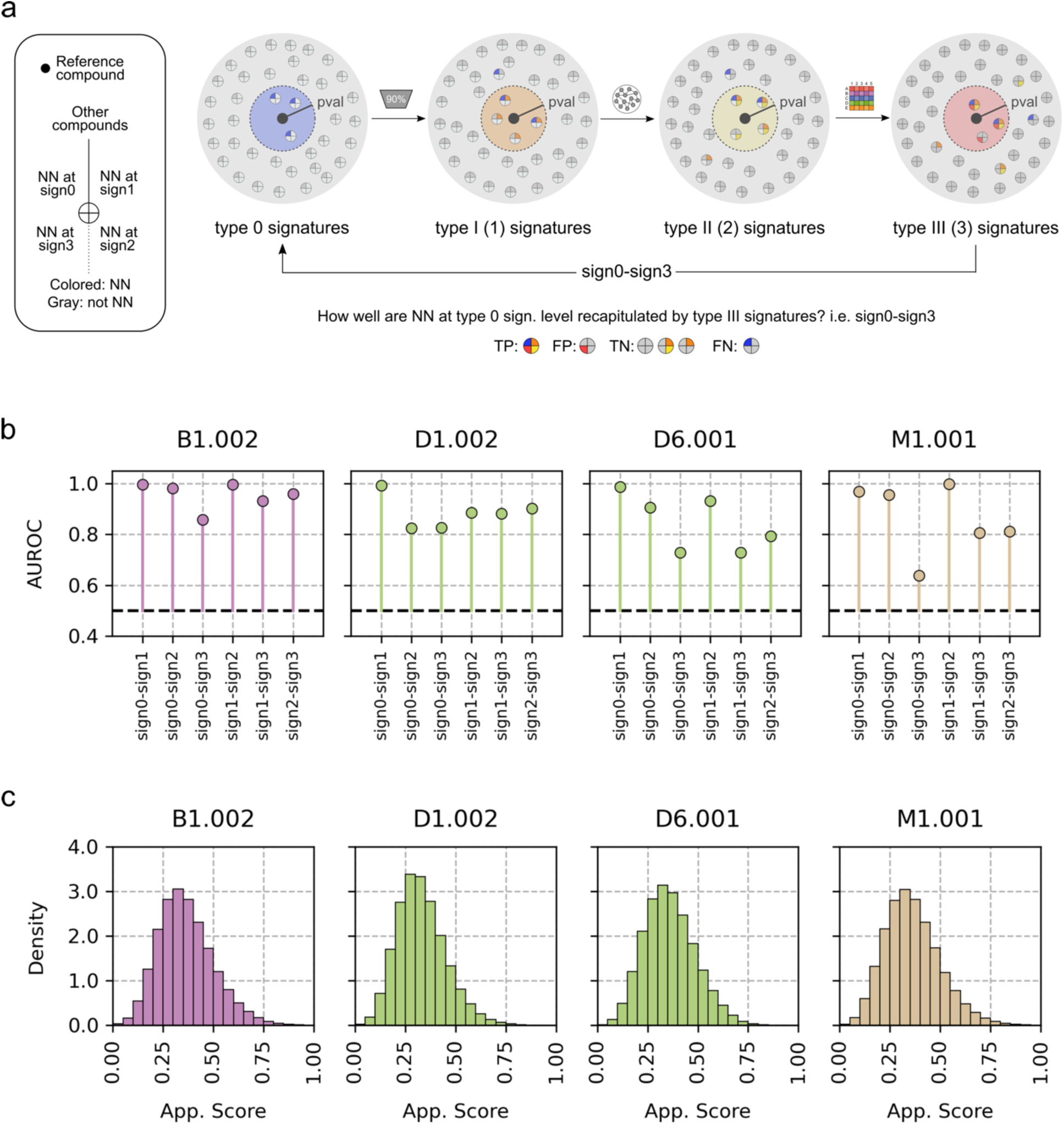
Quantitative evaluation of the signaturization process. **a)** Graphical representation of the validation strategy (from left to right). A reference compound (black circle) has a set of NN defined at type 0 signature level. Those NN may or may not be NN along the data compression, harmonization and integration pipeline. The better the ability to recapitulate NN defined at type 0 signature level by any other signature type, the fewer amount of information is lost along the computational pipeline. **b)** For each newly created CC space and pair of signature types (signA-signB), ability (y-axis, AUROC) from signBs to recapitulate NN defined at signA level. In brief, we defined a cutoff cosine distance for all signature types (0, I, II and III) at a pvalue of 0.01 (10,000 randomly selected compounds x 10 subsamples, considering the average value). Then, for each combination of signA-signB, we evaluated the ability of signB to recapitulate NN defined at signA level with a pvalue of 0.01 (for each combination, 2,500 random molecules, 5 subsamples; totalling 6 combinations of signature type pairs). **c)** For each newly created CC space, distribution of applicability values associated to the generated type III signatures.

Interestingly, we observed that, apart from including a higher number of compounds and MoAs, the new B1.002 space was less redundant than B1.001 (**Fig S1** and **Fig S2**). A tSNE 2D representation of these two spaces show how the incorporation of new data populates some areas of the target space not previously covered, and that the new B1.002 space was able to better recapitulate known MoAs and ATCs for annotated compounds (**Fig S3**).

A global picture of the pipeline with detailed steps, numbers and timings is shown in **Fig 2a**. Diagnosis plots for all signatures are shown in **Fig S1** and **Fig S2**.

B1.002 type 0, I, II and III signatures are available in https://zenodo.org/records/14000467.

### Task 2: Rebuilding a preexisting CC space with a different strategy of processing the raw data

#### Dataset description

Gene expression (GEx) responses after compound perturbation provide a convenient way to connect diseases to drugs. Since the release of the Connectivity Map (CMap) and LINCS resources, compound GEx signatures have allowed the identification of potential treatments for diverse diseases (e.g.^12,24,25^) and have provided new insights on compounds’ mechanisms of action (MoAs), compounds’ target and sensitivity responses. Given the significant value of these screenings, conscious efforts have been carried out to remove potential sources of noise and inherent biases, unearthing higher and more robust biological signals. Particularly relevant has been the implementation of the Characteristic Direction (chdir) method^26^ which, having been tailored in the context of compound perturbation responses, it is not only more suitable for detecting meaningful GEx changes but also provides a means to identify non-robust, noisy profiles. Thus, we re-processed raw LINCS data using the chdir GEx profile protocol, as indicated by the authors. Eventually, we ended up with 11,066 compounds (rows) and 12,328 genes (columns), which significantly enhanced the interpretability compared to the previous processing method. In the next paragraphs we exemplify the rebuilding of the D1 space with this new pre-processing protocol (D1.002).

#### Signaturization results ∼ 9 hours

After processing the data, we started the rebuilding of the D1 space (D1.002) with 11,066 compounds (rows) and 12,328 genes (columns), while our old processing (D1.001) contained 14,098 compounds and 7,967 CMap features. This reduction in the number of compounds is due to the additional chdir filtering step that accounts for the robustness in the GEx replicates. Now, each compound-gene pair was associated with a numerical value representing the induced gene expression change. After creating the new CC instance and loading the preprocessed bioactivity data, we obtained type 0 signatures for 11,066 compounds against 12,328 genes. It is important to note that, in the generation of type 0 signatures, we set the number of maximally considered features in the sanitization process up to 13 thousands, since the default value is 10 thousands (please see *Troubleshooting* and *Supplementary code: Task 2*). In addition, we observed that signatures were not redundant and that most values were in the [-0.05, 0.05] range (**Fig S4a**). Then, we obtained type I signatures for 11,065 compounds with 3,794 dimensions and generated harmonized type II signatures for these 11,065 compounds (**Fig S4b, c**). Finally, we integrated the D1.002 space with the rest of CC spaces and obtained type III signatures for all 1,009,305 compounds (**Fig S5**). To assess the capacity of type I, II and III signatures to recapitulate NN defined at type 0 signature level (raw bioactivity data), we implemented the methodology extensively described in the previous section (*Anticipated results: Task 1)* and illustrated in **Fig 3a**. In this task, we also observed that type III signatures were able to greatly recapitulate similarities defined at the raw bioactivity data level (type 0 signatures, AUROCs∼0.83, **Fig 3b**) and showed applicability values in the same range of preexisting CC bioactivity spaces (**Fig 3c**).

A downstream comparison with the space obtained from our old processing (D1.001) confirmed the overall improved consistency of this new space, as illustrated by the better recapitulation of known drug bioactivity associations, including their MoA and therapeutic (ATC) areas (**Fig S6**). This exercise highlights the importance not only of the quality of the raw data employed to develop the bioactivity descriptors but also of the preprocessing strategies.

A global picture of the pipeline with detailed steps, numbers and timings is shown in **Fig 2b**.

D1.002 type 0, I, II and III signatures are available in https://zenodo.org/records/14000494.

### Task 3: Creating a new CC space from a novel data type that fits the current CC universe

#### Dataset description

Deconvoluting the mechanisms of action (MoAs) of bioactive compounds within human cells is fundamental to tailor the eventual therapeutic effects they may exert. To study changes in protein levels induced by the presence of pharmacological agents, Mitchell et al.^14^ screened a collection of 875 small molecule perturbagens in HCT116 cells and derived the so-called ‘proteome fingerprints’ for them, accounting for the changes in protein abundance caused after 24h of treatment and achieving deep proteome coverage (>9,000 quantified proteins). In short, they observed that over half of the quantified proteome changed by at least twofold with at least one compound, and 98% of the proteins showed a variation of five standard deviations produced by at least one compound.

Although cell-based assays are indeed included in the CC (e.g. transcriptomics in D1 or sensitivity profiles in D2), proteomics data are not specifically considered. The main purpose of this exercise is to create a new CC space with the proteomics data presented above (namely D6.001) accounting for 875 compounds and 9,960 proteins.

#### Signaturization results ∼ 9 hours

After creating the new CC instance and loading the preprocessed bioactivity data, we found that 12.5% of the pairs did not have a reported (log2) fold change measurement (NaNs). We then obtained type 0 signatures for 873 compounds against 9,960 proteins and we observed that signatures were binary and completely non-redundant (**Fig S7a**). After that, we reduced their dimensionality by generating type I signatures for 873 compounds with 544 features and we then obtained type II signatures for these 873 compounds, at that point already harmonized with the other CC spaces (128 features). Finally, we derived type III signatures for these 873 compounds together with all the CC universe of small molecules, totaling to 1,009,363 signatures (**Fig S8**). To assess the capacity of type I, II and III signatures to recapitulate NN defined at type 0 signature level (raw bioactivity data), we implemented the methodology extensively described in the section *Anticipated results: Task 1* and illustrated in **Fig 3a**. In brief, we observed that type III signatures were mostly able to recapitulate similarities defined at the raw bioactivity data level (type 0 signatures, AUROCs∼0.73, **Fig 3b**), although the performance was markedly lower than in other new spaces (i.e. B1.002 and D1.002). In addition, we also observed that applicability values associated to type III signatures where in the same numerical range than in preexisting CC bioactivity spaces (**Fig 3c**).

Additionally, when we compared the proteomics signatures to those derived from transcriptomics, we observed that proteomics-based D6.001 type III signatures were able to recapitulate known MoAs at pair with the new D1.002 signatures, and better that the original D1.001 space (**Fig S9a, c, e**). D6.001 is also better at capturing therapeutic areas (ATC) than the transcriptional spaces, perhaps reflecting protein abundance is a closer proxy for clinical outcomes (**Fig S9b, d, f**). We also asked if proteomics-derived signatures were able to recapitulate nearest-neighbors in transcriptomics signatures, finding a moderate capacity (AUROC of 0.72 and 0.74 for D1.001 and D1.002, respectively), which indicates a high degree of complementarity between the two bioactivity spaces (**Fig S9g, h**).

A global picture of the pipeline with detailed steps, numbers and timings is shown in **Fig 2c**. Diagnosis plots for all signatures are shown in **Fig S7** and **Fig S8**.

D6.001 type 0, I, II and III signatures are available in https://zenodo.org/records/14000554.

### Task 4: Creating a new CC space from a novel data type that does not fit the current CC universe

#### Dataset description

Non-antibiotic marketed drugs have been recently associated with changes in the composition of the human gut microbiome^27^. Such unforeseen activity may explain potential antibiotic-like side effects, might promote antibiotic resistance and may eventually contribute to a significant decrease in microbiome diversity^28^. To unravel potential relationships between non-antibiotic drugs and microbes, Maier et al.^28^ screened a collection of drug-like compounds (including human-targeted drugs, antibiotics, anti-infective compounds and veterinary drugs, among others) against a set of 40 representative gut bacterial strains. Indeed, they observed that 24% of the drugs having human targets were able to inhibit the growth of at least one gut bacterial strain *in vitro*. We exhaustively processed their data and defined a binary growth inhibition matrix between tested compounds and strains. The novel bioactivity data involved 1,155 compounds (marketed drugs, rows) tested against 40 representative gut bacterial strains (columns). The main purpose of this exercise is to create a new CC space (i.e. M1.001) where the chemical compounds exert their bioactivities on microbial species in the human microbiota.

#### Signaturization results ∼ 5 hours

After creating the new CC instance and loading the preprocessed bioactivity data, we obtained type 0 signatures for 310 compounds against 40 gut bacterial strains. In fact, we found that 769 molecules exhibited null bioactivity profiles and were thus filtered out from the signaturization process (*min_keys_abs* parameter). Additionally, 76 compounds were removed due to an excessive proportion of positive features (*max_keys_freq* parameter). After that, we obtained type I signatures for 200 compounds against 13 features, significantly reducing the number of dimensions of the data (**Fig S10**). We then generated type II signatures for these molecules, harmonizing the number of features to all the other spaces in the CC (128 features). Finally, we derived type III signatures for these 200 compounds and for all the remaining molecules in the CC universe (1,009,305 molecules, in total). To assess the capacity of type I, II and III signatures to recapitulate NN defined at type 0 signature level (raw bioactivity data), we implemented the methodology extensively described in the section *Anticipated results: Task 1* and further illustrated in **Fig 3a**. And, again, we observed a decreased capacity for type III signatures to recapitulate similarities defined at the raw bioactivity data (type 0 signatures, AUROCs∼0.64, **Fig 3b**) compared with the three previous examples (i.e. B1.002, D1.002 and D6.001). This lower performance highlighted the very orthogonal nature of the bioactivities in the M space (i.e. effect of molecules on microbiota) and the rest of the CC, clearly dominated by human data. In addition, we also observed that applicability values associated to type III signatures where in the same numerical range than in preexisting CC bioactivity spaces (**Fig 3c**).

The construction of the M1.001 space was challenging since, despite the apparent initially high number of compounds and features (i.e. microbial species), the final number or compound-microbe relationships was quite limited. Nevertheless, we still observed a remarkable capacity of type III signatures to recapitulate known MoAs and ATC for annotated compounds (**Fig S11**).

A global picture of the pipeline with detailed steps, numbers and timings is shown in **Fig 2d**. Diagnosis plots for all signatures are shown in **Fig S10** and **S11**.

M1.001 type 0, I, II and III signatures are available in https://zenodo.org/records/14000572.

#### Troubleshooting

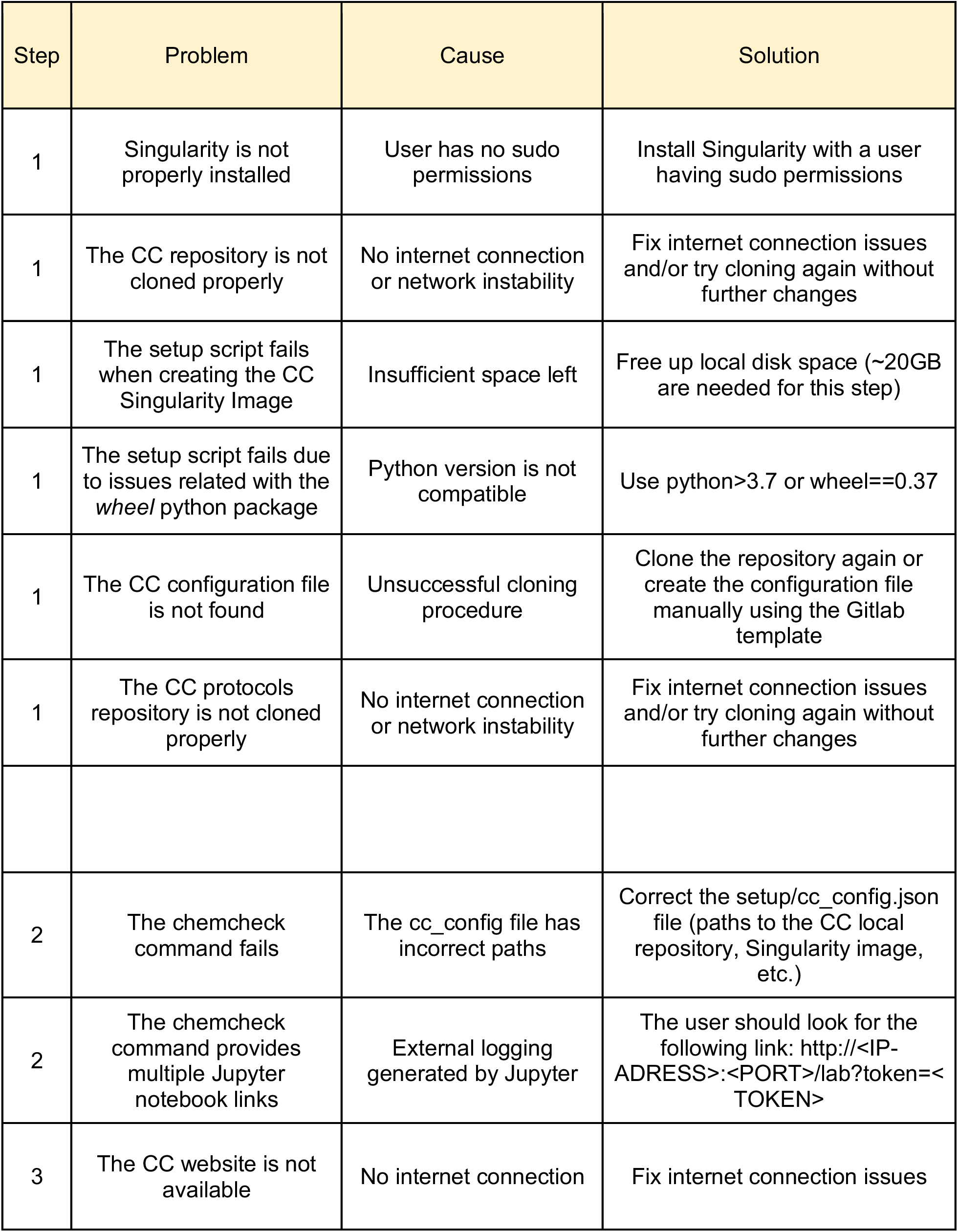
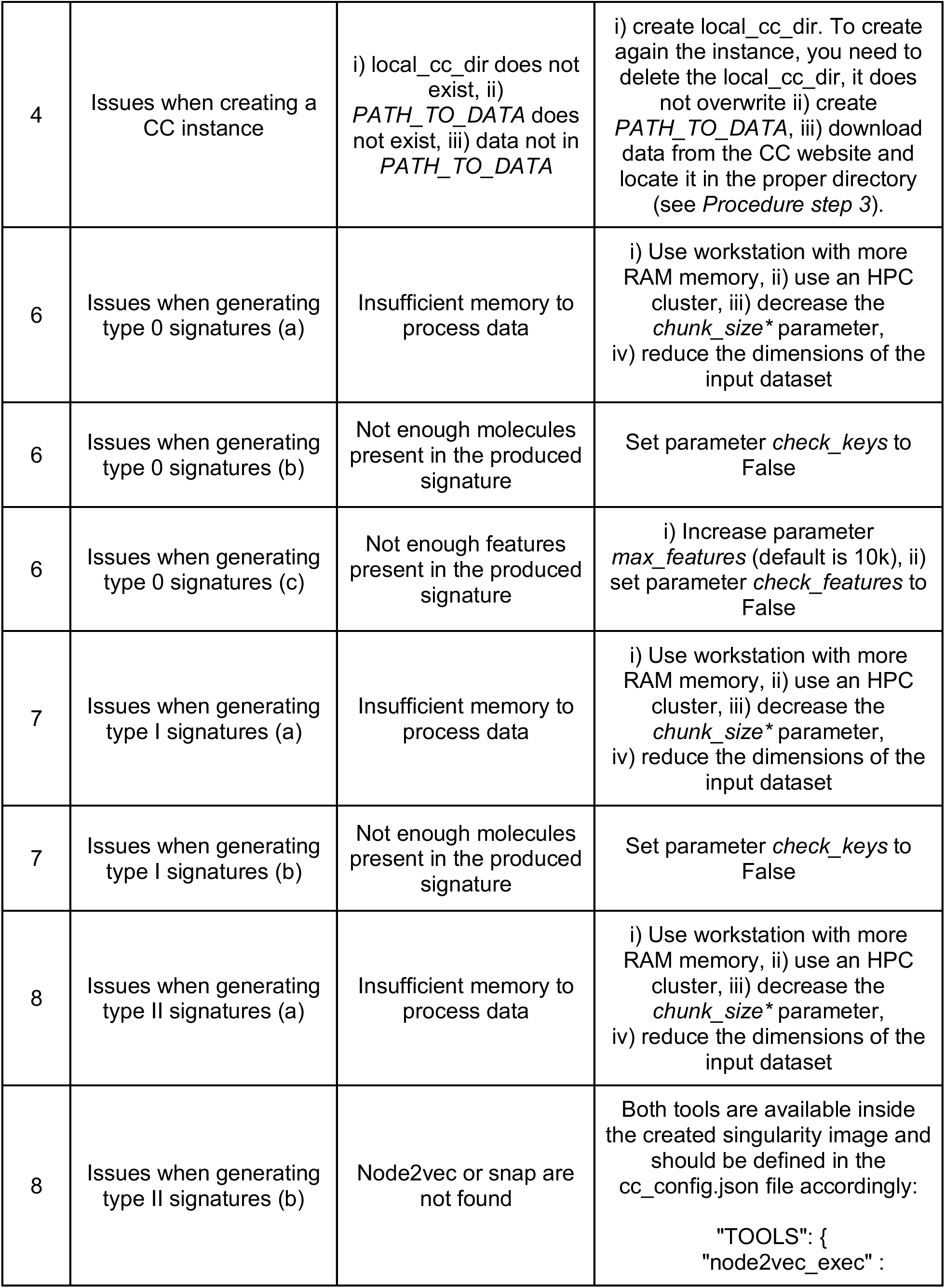
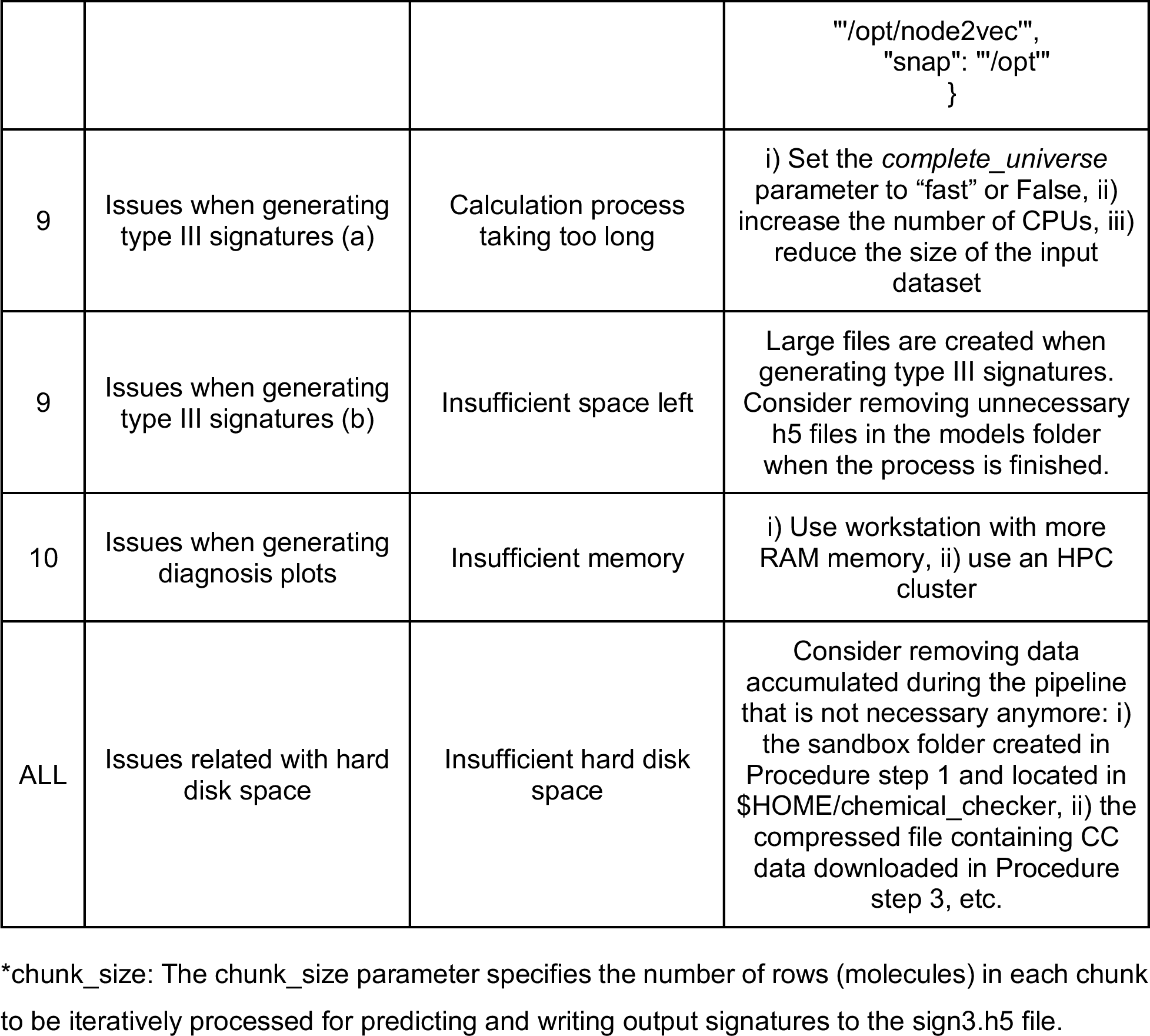

| Step | Problem | Cause | Solution |
| --- | --- | --- | --- |
| 1 | Singularity is not properly installed | User has no sudo permissions | Install Singularity with a user having sudo permissions |
| 1 | The CC repository is not cloned properly | No internet connection or network instability | Fix internet connection issues and/or try cloning again without further changes |
| 1 | The setup script fails when creating the CC Singularity Image | Insufficient space left | Free up local disk space (~20GB are needed for this step) |
| 1 | The setup script fails due to issues related with the *wheel* python package | Python version is not compatible | Use python>3.7 or wheel==0.37 |
| 1 | The CC configuration file is not found | Unsuccessful cloning procedure | Clone the repository again or create the configuration file manually using the Gitlab template |
| 1 | The CC protocols repository is not cloned properly | No internet connection or network instability | Fix internet connection issues and/or try cloning again without further changes |
| 2 | The chemcheck command fails | The cc_config file has incorrect paths | Correct the setup/cc_config.json file (paths to the CC local repository, Singularity image, etc.) |
| 2 | The chemcheck command provides multiple Jupyter notebook links | External logging generated by Jupyter | The user should look for the following link: http://<IP-ADRESS>:<PORT>/lab?token=<TOKEN> |
| 3 | The CC website is not available | No internet connection | Fix internet connection issues |
| 4 | Issues when creating a CC instance | i) `local_cc_dir` does not exist, ii) `PATH_TO_DATA` does not exist, iii) data not in `PATH_TO_DATA` | i) create `local_cc_dir`. To create again the instance, you need to delete the `local_cc_dir`, it does not overwrite ii) create `PATH_TO_DATA`, iii) download data from the CC website and locate it in the proper directory (see *Procedure step 3*). |
| 6 | Issues when generating type 0 signatures (a) | Insufficient memory to process data | i) Use workstation with more RAM memory, ii) use an HPC cluster, iii) decrease the `chunk_size*` parameter, iv) reduce the dimensions of the input dataset |
| 6 | Issues when generating type 0 signatures (b) | Not enough molecules present in the produced signature | Set parameter `check_keys` to False |
| 6 | Issues when generating type 0 signatures (c) | Not enough features present in the produced signature | i) Increase parameter `max_features` (default is 10k), ii) set parameter `check_features` to False |
| 7 | Issues when generating type I signatures (a) | Insufficient memory to process data | i) Use workstation with more RAM memory, ii) use an HPC cluster, iii) decrease the `chunk_size*` parameter, iv) reduce the dimensions of the input dataset |
| 7 | Issues when generating type I signatures (b) | Not enough molecules present in the produced signature | Set parameter `check_keys` to False |
| 8 | Issues when generating type II signatures (a) | Insufficient memory to process data | i) Use workstation with more RAM memory, ii) use an HPC cluster, iii) decrease the `chunk_size*` parameter, iv) reduce the dimensions of the input dataset |
| 8 | Issues when generating type II signatures (b) | Node2vec or snap are not found | Both tools are available inside the created singularity image and should be defined in the `cc_config.json` file accordingly:<br>`"TOOLS": { "node2vec_exec" :` |
|  |  |  | `"/opt/node2vec", "snap": "/opt" }` |
| 9 | Issues when generating type III signatures (a) | Calculation process taking too long | i) Set the *complete_universe* parameter to "fast" or False, ii) increase the number of CPUs, iii) reduce the size of the input dataset |
| 9 | Issues when generating type III signatures (b) | Insufficient space left | Large files are created when generating type III signatures. Consider removing unnecessary h5 files in the models folder when the process is finished. |
| 10 | Issues when generating diagnosis plots | Insufficient memory | i) Use workstation with more RAM memory, ii) use an HPC cluster |
| ALL | Issues related with hard disk space | Insufficient hard disk space | Consider removing data accumulated during the pipeline that is not necessary anymore: i) the sandbox folder created in Procedure step 1 and located in \$HOME/chemical_checker, ii) the compressed file containing CC data downloaded in Procedure step 3, etc. |

\*chunk\_size: The chunk\_size parameter specifies the number of rows (molecules) in each chunk to be iteratively processed for predicting and writing output signatures to the sign3.h5 file.

#### Timing

| Step(s)* | Description | Timing |
| --- | --- | --- |
| 1 | Installing the CC | 30 min |
| 2 | Opening a Jupyter Notebook within the CC image | 10 sec |
| 3 | Gathering CC data | 45 min |
| 4 [1, 2, 3, 4] | Creating a CC instance | 10 / 10 / 10 / 10 sec |
| 5 [1, 2, 3, 4] | Loading raw data | 10 / 10 / 10 / 10 sec |
| 6 [1, 2, 3, 4] | Generating type 0 signatures | 1 / 5 / 1 / 1 min |
| 7 [1, 2, 3, 4] | Generating type I signatures | < 10 / 30 / 10 / 10 min |
| 8 [1, 2, 3, 4] | Generating type II signatures | 2 / 10 / 1 / 1 min |
| 9 [1, 2, 3, 4] | Generating type III signatures** | 8 / 10 / 10 / 6 h |
| 9 [1, 2, 3, 4] | Generating type III signatures*** | 7 / 8 / 8 / 4 h |
| 10 | Generating diagnosis plots (small) | < 2 min each |
| 10 | Generating diagnosis plots (medium) | ~ 45 min each |
| 4-9 [1] | Task 1: B1.002 | ~ 9h** / 8h*** |
| 4-9 [2] | Task 2: D1.002 | ~ 11h** / 9h*** |
| 4-9 [3] | Task 3: D6.001 | ~ 11h** / 9h*** |
| 4-9 [4] | Task 4: M1.001 | ~ 7h** / 5h*** |

\*Step nomenclature: Step [Task]
\*\*The generation of type III signatures was run in a 32-cores CPU machine
\*\*\*The generation of type III signatures was run in a 12-cores GPU machine.

## Conclusions

Chemical signatures encode the physicochemical and structural properties of small molecules in the form of numerical descriptors, and they are at the core of chemical comparisons and search algorithms^29^. Recently, the widespread availability of bioactivity data has led to enhanced compound representations that capture their biological effects along with their chemical structures. Yet, such bioactivity descriptors are scarce, limited mostly to a small number of well-documented molecules. To overcome this problem, we implemented a collection of deep neural networks able to leverage the experimentally determined bioactivity data associated to small molecules and infer the missing bioactivity signatures for any compound of interest^10^. Originally, our strategy related to bioactivities of 25 different types (including target profiles, cellular responses and clinical outcomes), as described in the Chemical Checker publication^9^. However, unlike typical chemical descriptors that remain static, our bioactivity signatures are dynamic, evolving as new bioactivity data accumulates in databases or new strategies to process it appear.

We have now made available the complete computational protocol to modify or generate novel bioactivity spaces and signatures, describing the main steps needed to leverage diverse bioactivity data with the Chemical Checker using the predefined data curation pipeline. Moreover, we have illustrated the functioning of the protocol through four specific examples, including the incorporation of new compounds to an already existing bioactivity space, a change in the data pre-processing without altering the underlying experimental data, and the creation of two novel bioactivity spaces from scratch.

Despite the regular updates, the expansion of the CC resource, beyond the original 25 spaces, is constrained by the availability and quality of public data sets. However, the systematic measurement of drug-induced perturbations in biological systems through omics technologies, together with advancements in phenotypic screening, are becoming increasingly common in the public and private sectors, providing an ever-growing corpus of biomolecular information. Besides, given the vast diversity of drug-like small molecules and the consolidation of generative AI as a novel strategy to design new chemical entities, computational tools are progressively becoming essential for initially estimating the biological effects of compounds. Overall, we envision that this protocol to create new bioactivity spaces and the associated descriptors will serve as a pivotal tool for exploring the bioactivity spectrum of compounds, effectively bridging the existing knowledge gap between the chemical structure and the exerted biological effects of small molecules.

## Author contributions statements

M.B., M. D-F., O.G-P. and P.A. designed the study. M.B., M.D-F., O.G-P., M.L., Y.M. and N.S. implemented the computational pipeline. A.C-C., A. F-T., N.K., E.P-L., G.R-G., and E.V. created the different novel bioactivity spaces. A.C-C. and P.A. wrote the manuscript. All authors analyzed the results, read and approved the manuscript.

## Acknowledgments

P.A. acknowledges the support of the Generalitat de Catalunya (2021 SGR 00876), the Spanish Ministerio de Ciencia, Innovación y Universidades (PID2020-119535RB-I00), the Instituto de Salud Carlos III (IMPaCT-Data), and the European Commission (CLARITY: 101137201). A.C-C. and E.P-L are recipients of an FI fellowship (2020 FI_B 00094 and 2022 FI_B 00767). N.K. was supported by a European Molecular Biology Organization postdoctoral fellowship (ALTF 906-2021).

