## Supplementary Information for "Integration of diverse bioactivity data into the Chemical Checker compound universe"

#### *Downloading the CC bioactivity data*

To implement all CC functionalities illustrated in this work, we recommend gathering the CC bioactivity data from our web servers using code provided *(see Procedure 2*). Since the CC undergoes annual updates, the size of the downloaded files may increase over time, along with the amount of publicly released bioactivity data. To partially overcome long download times or the need for extensive local storage, users may select specific files to download depending on the CC functionality they wish to use. In the following points, we provide a detailed overview of which CC functionalities are related to specific downloadable files, helping users decide what to download.

**-Protocols Functionalities Essentials:**

- **Full Type 0 Signatures for Chemical Spaces (A1-5):** Required for the complete universe functionality.
- **Reference Type I Signatures for Chemical Spaces (A1-5):** Required for the complete universe functionality.
- **Reference Type I Models for Chemical Spaces (A1-5):** Required for the complete universe functionality.
- **Reference Type II Signatures for Chemical Spaces (A1-5):** Required for the complete universe functionality.
- **Reference Type II Models for Chemical Spaces (A1-5):** Required for the complete universe functionality.
- **Full Type II Signatures:** Required for generating Type III signatures in a newly created CC space.
- **Full Type III Signatures:** Required for creating medium diagnosis plots for newly generated Type III signatures
- **Metadata:** Required for the generation of medium diagnosis plots.

**-Additional Signatures:**

Full Type 0, I or II signatures are necessary for generating medium diagnosis plots for newly created Type 0, I or II signatures. For this protocol, small diagnosis plots are provided for these signature types.

If users want to evaluate the recapitulation of the raw CC bioactivity data (i.e. type 0 signatures) by means of generated signatures (e.g. type III signatures of a newly created space), they will need to download Full type 0 signatures for all CC spaces (+20GB) and adjust the diagnosis plot code accordingly (i.e. *ref_cctype='sign0'*).

**-Reduced Data Version for users.**

For users not interested in the complete universe functionality, only **the Full Type II Signatures** (for creating Type III signatures) and the **Full Type III Signatures** (for diagnosis) are necessary.

##### *Implementing GPU computation*

In order for the singularity image to be able to use a GPU, the user needs to install the appropriate cuda libraries (cuda-toolkit and cudnn) and make sure they are visible from the image. We recommend creating a new conda environment locally with the same python version (3.10) and tensorflow version (2.8) as in the image and install the cuda libraries there. After that, the user must add the following line at the beginning of the *run_chemicalchecker.sh* file, pointing to the directories in which cuda is installed:

```
export SINGULARITYENV_LD_LIBRARY_PATH=<PATH_TO_MINICONDA>/miniconda3/pkgs/cudatoolkit-11.8.0-h6a678d5_0/lib:<PATH_TO_MINICONDA>/miniconda3/pkgs/cudnn-8.9.2.26-cuda11_0/lib
```

An alternative option, but more time consuming, is to install the cuda libraries in the singularity sandbox and generate the image again. The user can check whether the GPU is detected with the following line of code in the *chemcheck* notebook:

```
import tensorflow as tf
tf.config.list_physical_devices('GPU')
```

**-Installing cuda libraries:**

```
conda install nvidia::cuda-toolkit
conda install conda-forge::cudnn
```

#### Supplementary Figures

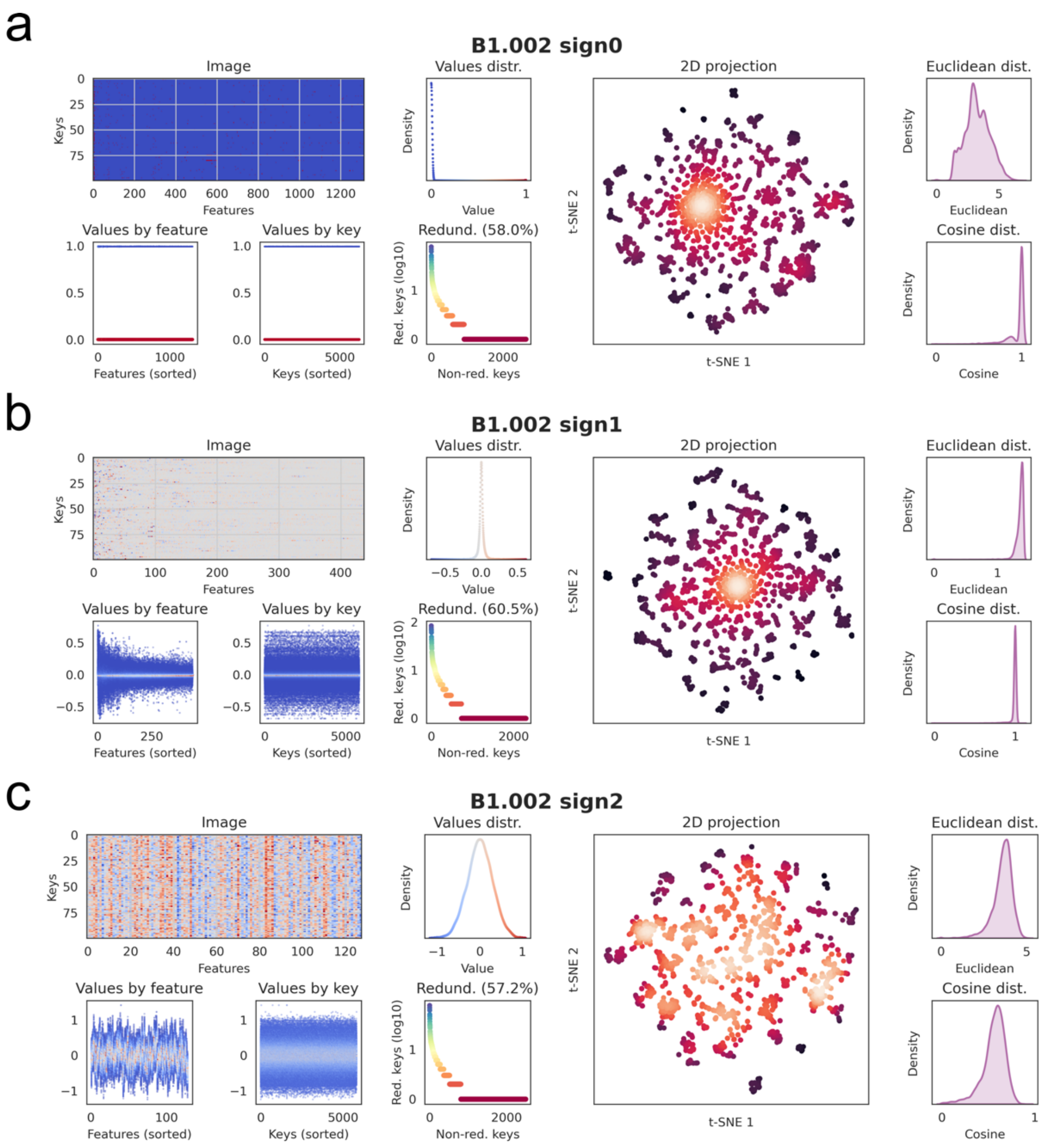


**Figure S1**: Diagnosis plots for the B1.002 space (please see *Anticipated Results: Task 1*). **a)** type 0 signatures, **b)** type I signatures and **c)** type II signatures. For further information about diagnosis plots, please see *Overview of the Chemical Checker data integration pipeline - Interpreting the results: diagnosis plots*.

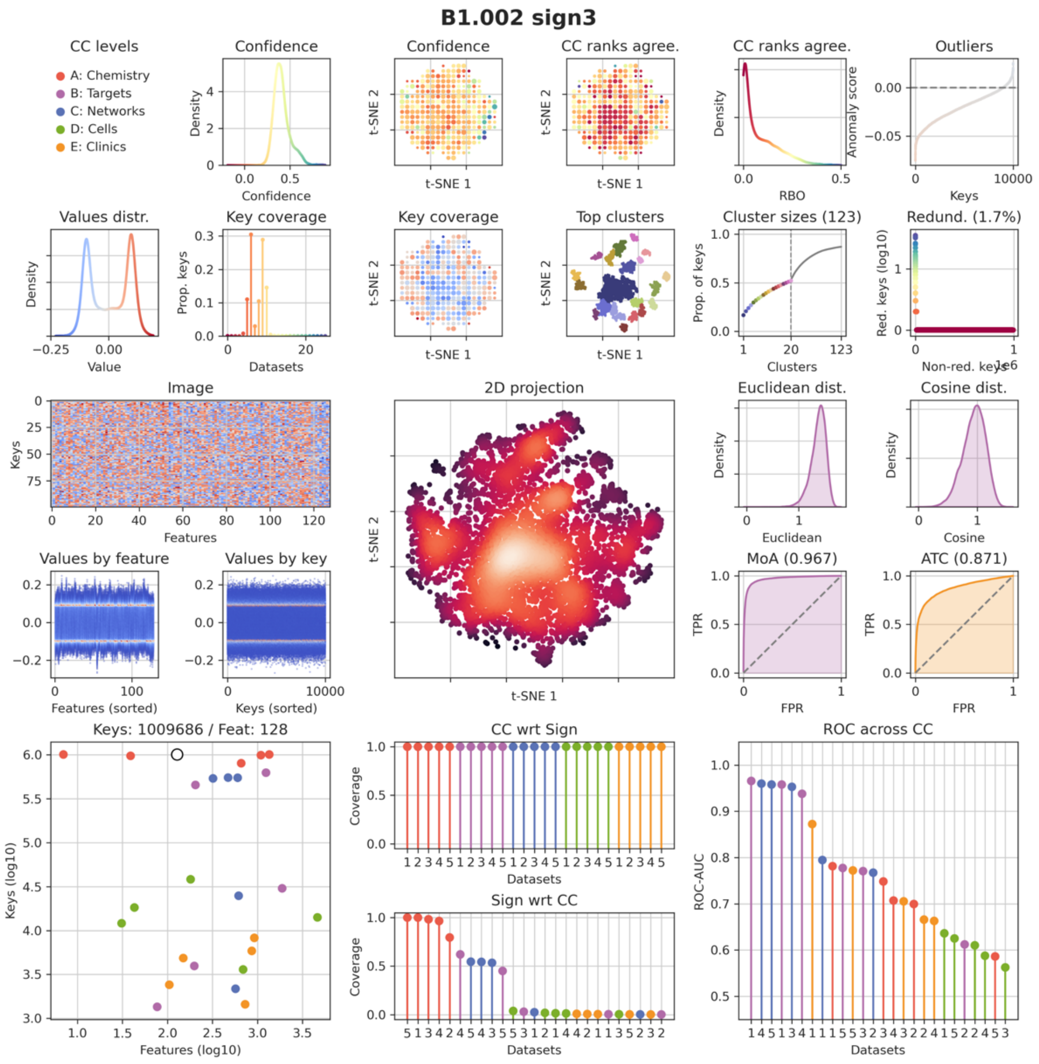


**Figure S2**: Extended diagnosis plots for B1.002 type III signatures (please see *Anticipated Results: Task 1*). For further information about diagnosis plots, please see *Overview of the Chemical Checker data integration pipeline - Interpreting the results: diagnosis plots* and check our Gitlab repository.

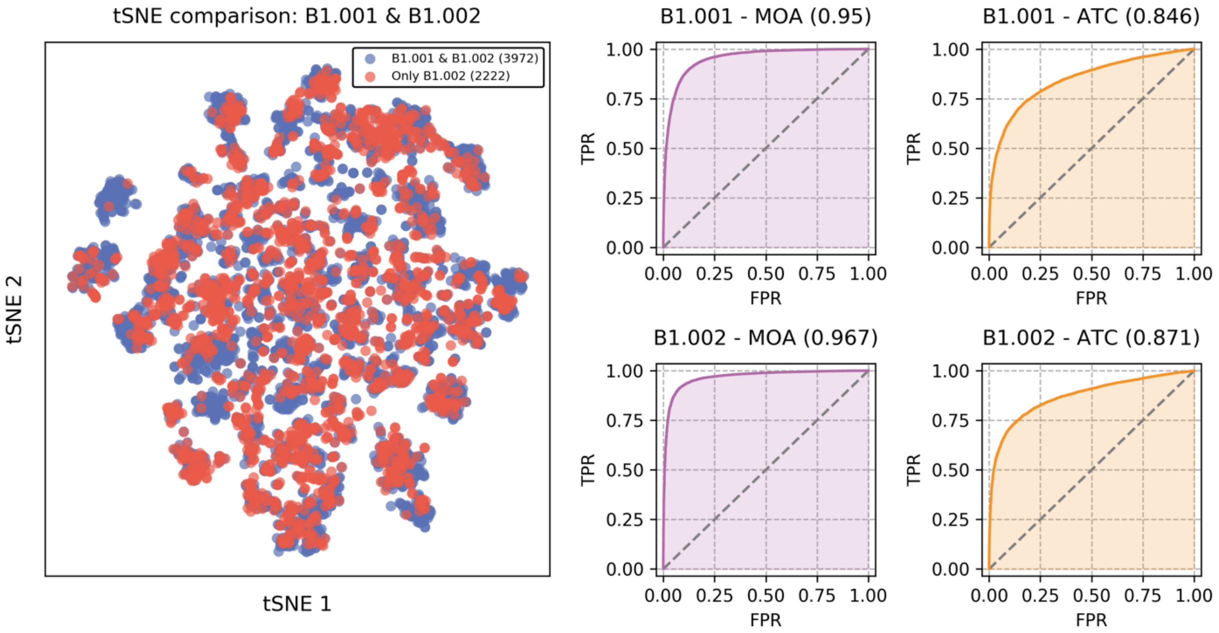


**Figure S3**: Comparison between B1.001 and B1.002. Left: 2D tSNE representation of B1.001 and B1.002 type III signatures having a corresponding type 0 signature in B1.001 and B1.002, respectively. Right: recapitulation of MOA (B1.001 type 0 signatures, purple) and ATC (E1.001 type 0 signatures, orange) using B1.001 (top) and B1.002 (bottom) type III signatures.

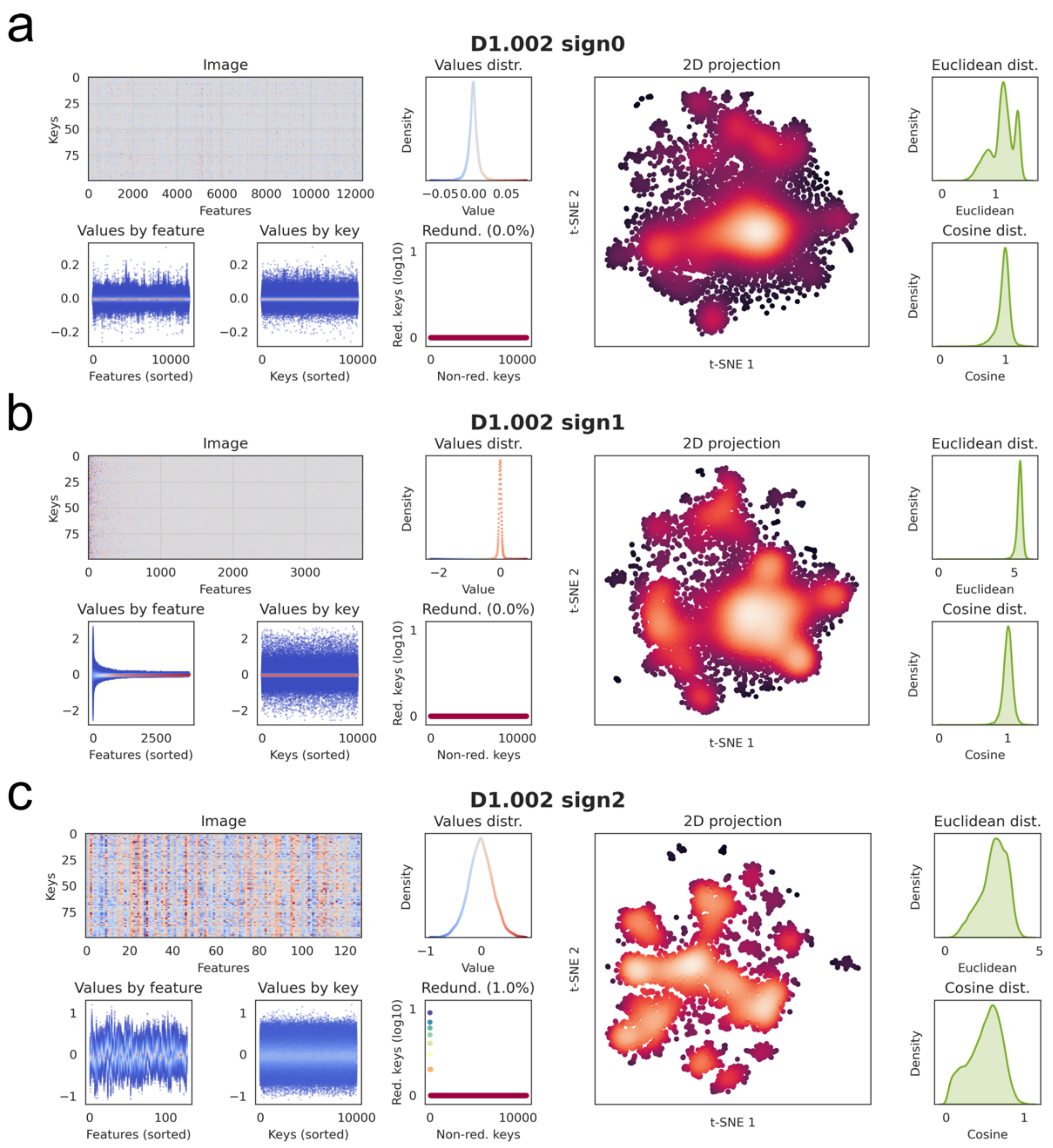


**Figure S4**: Diagnosis plots for the D1.002 space (please see *Anticipated Results: Task 2*). **a)** type 0 signatures, **b)** type I signatures and **c)** type II signatures. For further information about diagnosis plots, please see *Overview of the Chemical Checker data integration pipeline - Interpreting the results: diagnosis plots*.

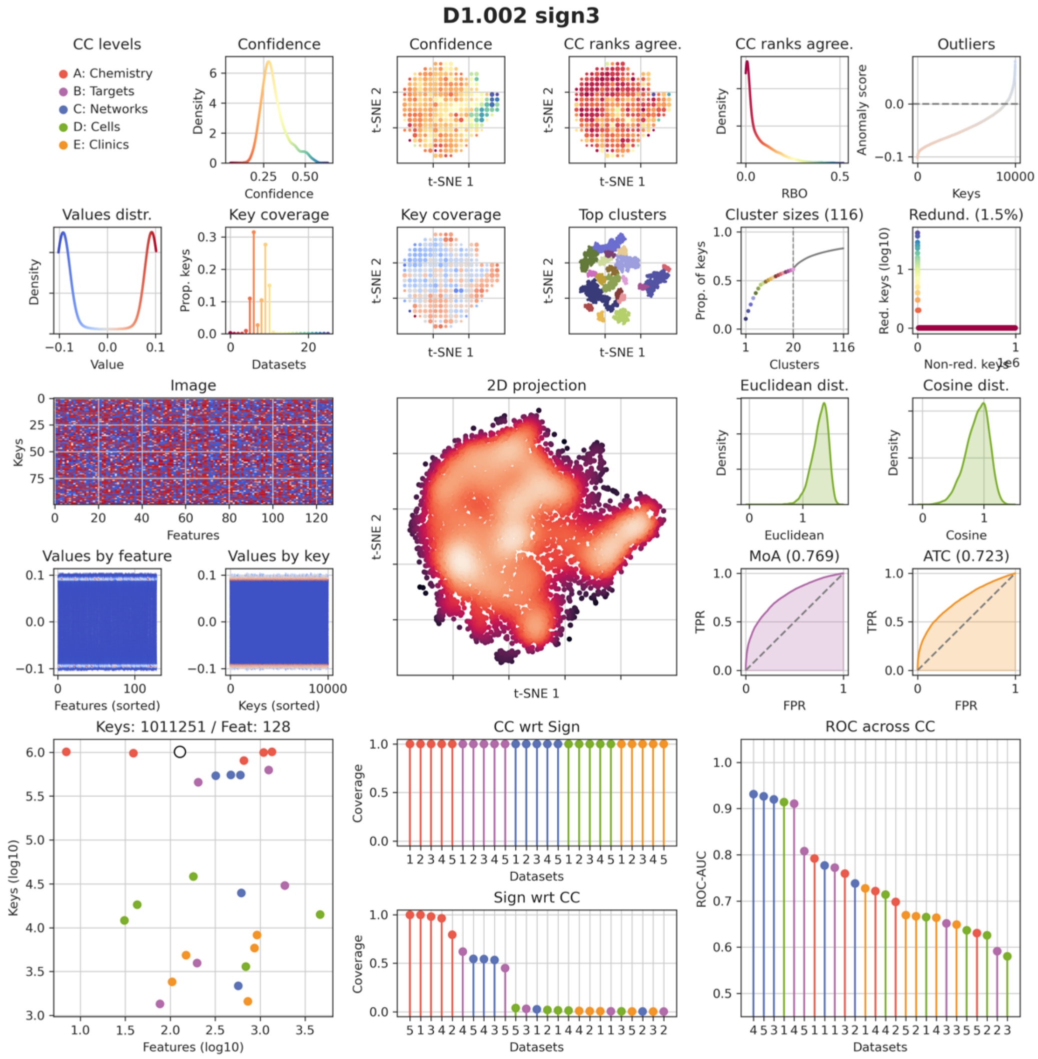


**Figure S5**: Extended diagnosis plots for D1.002 type III signatures (please see *Anticipated Results: Task 2*). For further information about diagnosis plots, please see *Overview of the Chemical Checker data integration pipeline - Interpreting the results: diagnosis plots* and check our Gitlab repository.

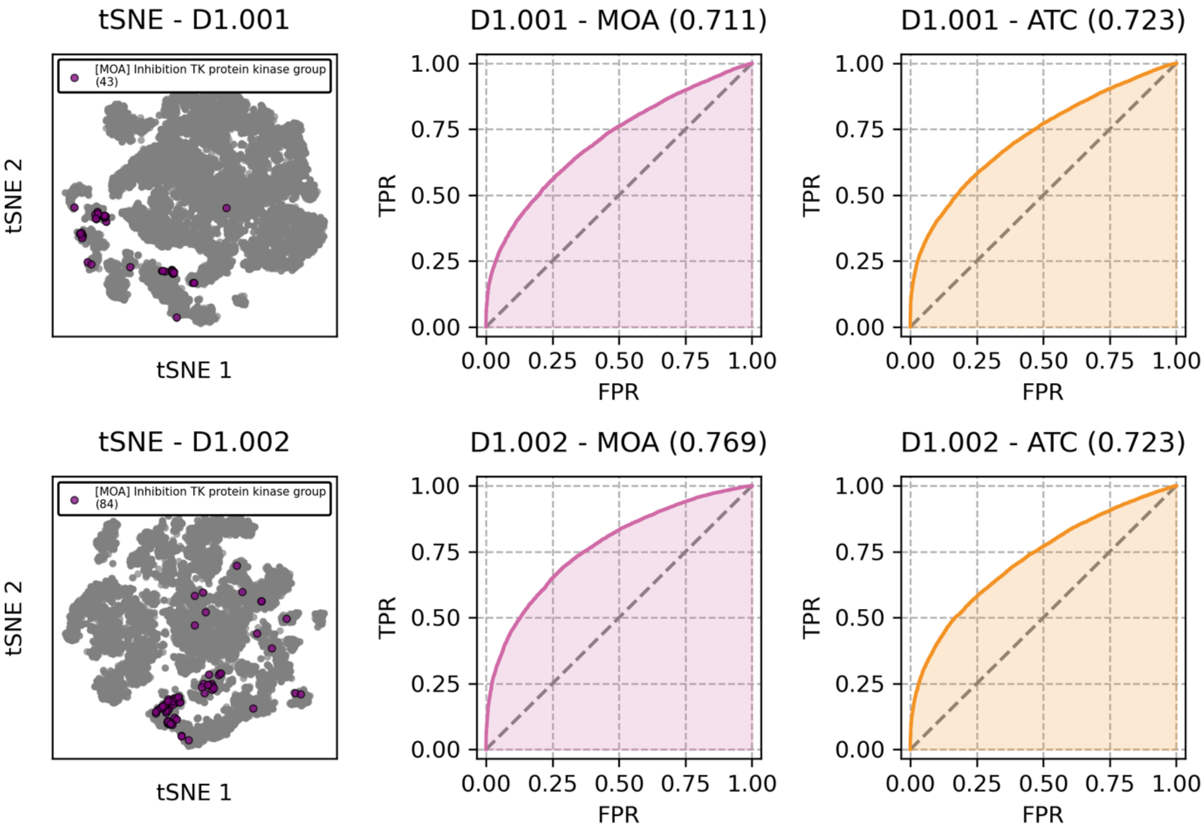


**Figure S6**: Comparison between D1.001 and D1.002. Left: 2D tSNE representation of D1.001 (top) and D1.002 (bottom) type III signatures having a corresponding type 0 signature in D1.001 and D1.002, respectively. Points highlighted in purple correspond to compounds related to the inhibition of the TK protein group. Center and right: recapitulation of MOA (B1.001 type 0 signatures, purple) and ATC (E1.001 type 0 signatures, orange) using D1.001 (top) and D1.002 (bottom) type III signatures.

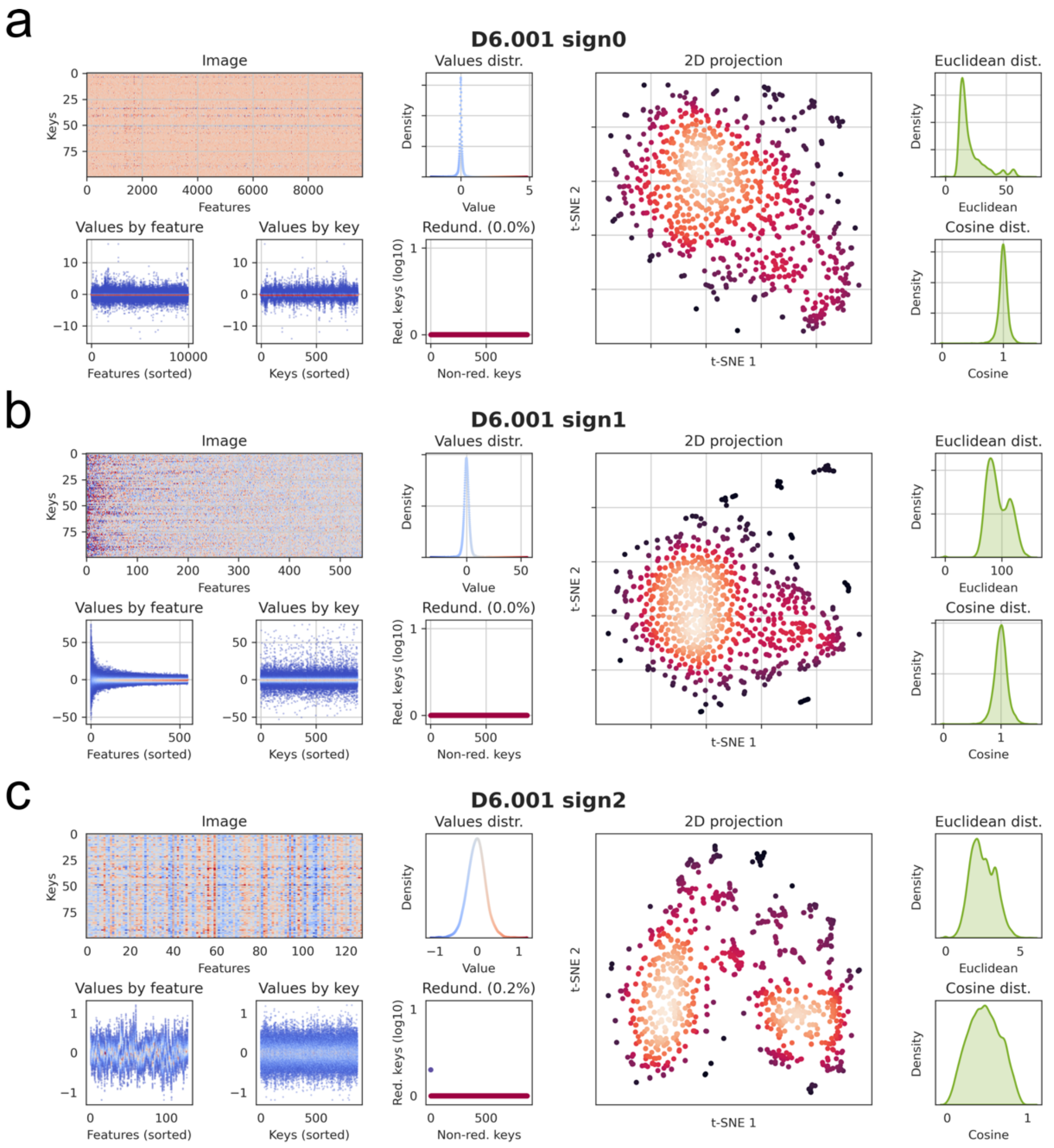


**Figure S7**: Diagnosis plots for the D6.002 space (please see *Anticipated Results: Task 3*). **a)** type 0 signatures, **b)** type I signatures and **c)** type II signatures. For further information about diagnosis plots, please see *Overview of the Chemical Checker data integration pipeline - Interpreting the results: diagnosis plots*.

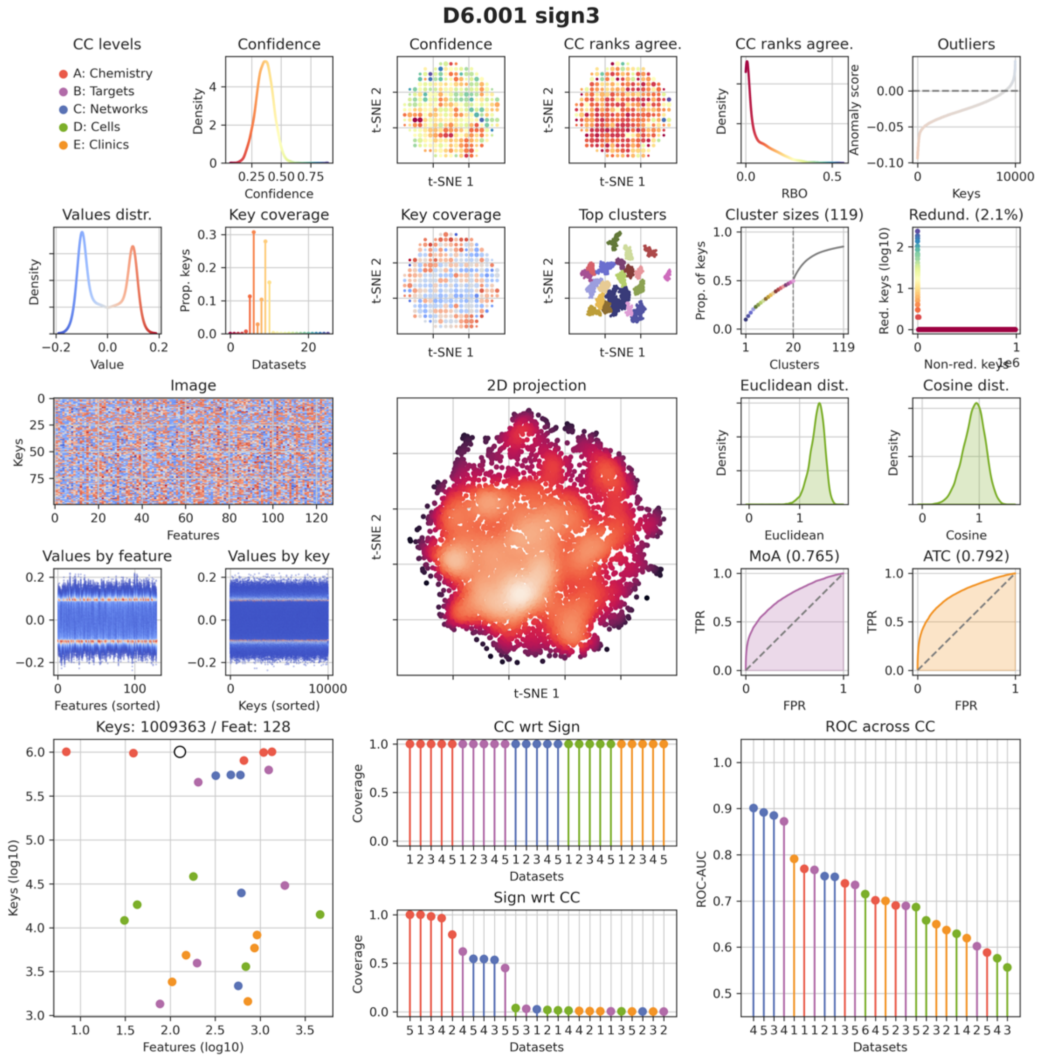


**Figure S8**: Extended diagnosis plots for D6.002 type III signatures (please see *Anticipated Results: Task 3*). For further information about diagnosis plots, please see *Overview of the Chemical Checker data integration pipeline - Interpreting the results: diagnosis plots* and check our Gitlab repository.

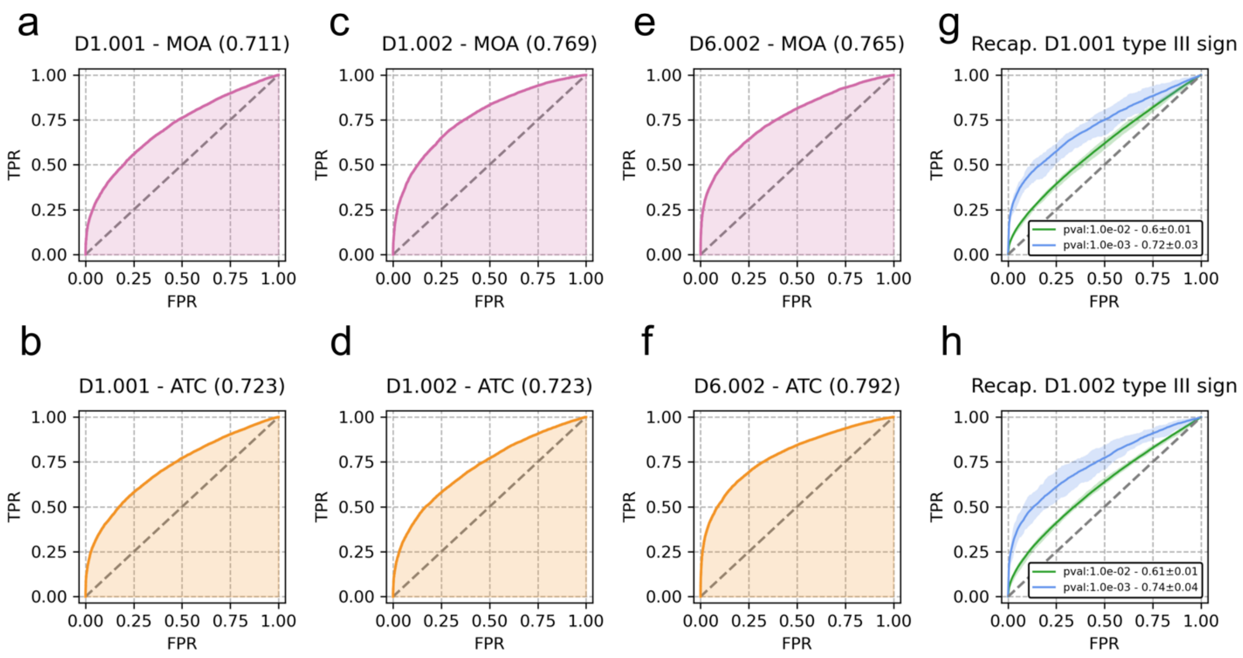


**Figure S9**: Comparison between D1.001, D1.002 and D6.001. **a)** Recapitulation of MoA (B1.001 type 0 signatures, purple) using D1.001 type III signatures. **b)** Recapitulation of ATC (E1.001 type 0 signatures, orange) using D1.001 type III signatures. **c)** Recapitulation of MoA (B1.001 type 0 signatures, purple) using D1.002 type III signatures. **d)** Recapitulation of ATC (E1.001 type 0 signatures, orange) using D1.002 type III signatures. **e)** Recapitulation of MoA (B1.001 type 0 signatures, purple) using D6.001 type III signatures. **f)** Recapitulation of ATC (E1.001 type 0 signatures, orange) using D6.001 type III signatures. **g)** Recapitulation of kNN at D1.001 type III signature level using D6.001 type III signatures at p-values 0.01 and 0.001. **h)** Recapitulation of NN at D1.002 type III signature level using D6.001 type III signatures at p-values 0.01 and 0.001.

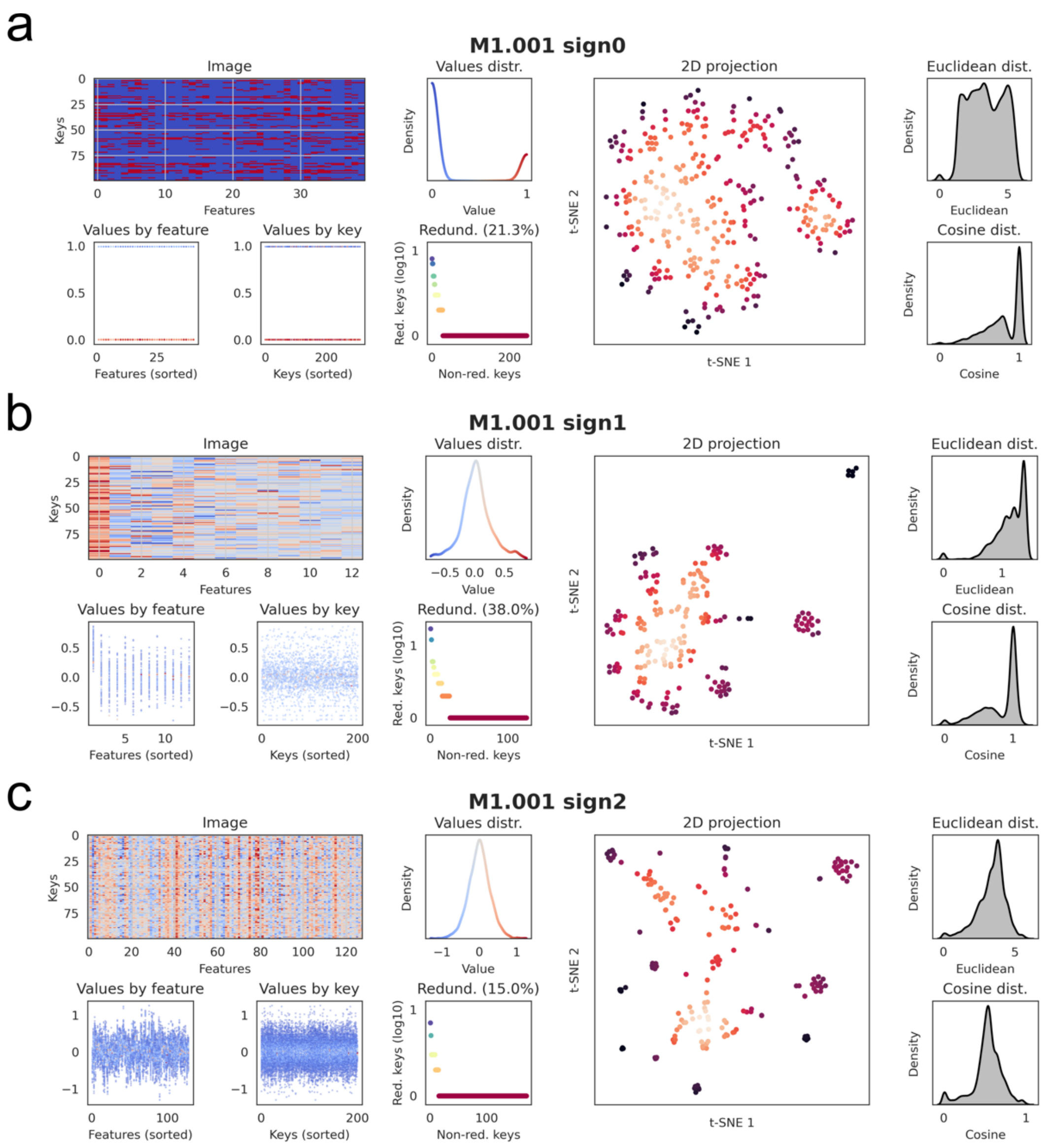


**Figure S10**: Diagnosis plots for the M1.001 space (please see *Anticipated Results: Task 4*). **a)** type 0 signatures, **b)** type I signatures and **c)** type II signatures. For further information about diagnosis plots, please see *Overview of the Chemical Checker data integration pipeline - Interpreting the results: diagnosis plots*.

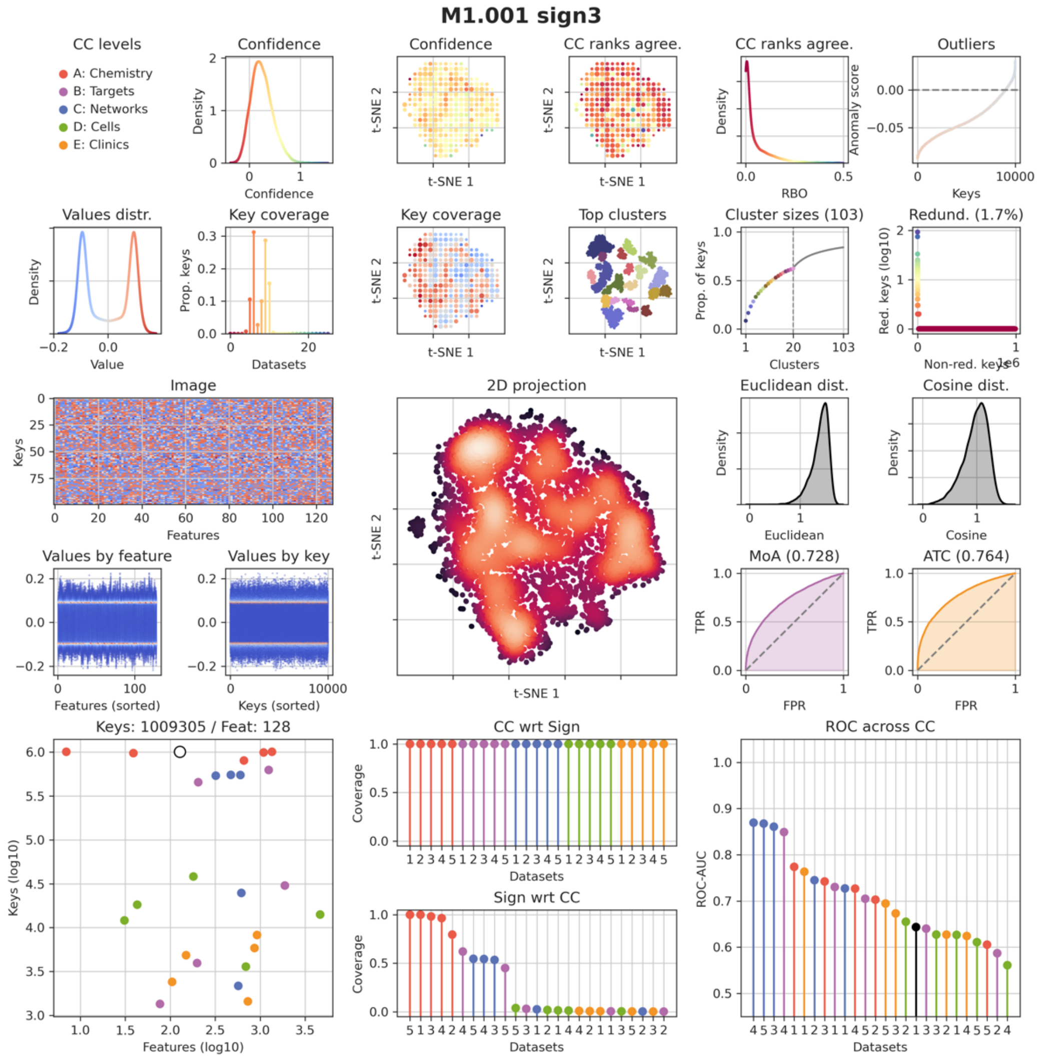


**Figure S11**: Extended diagnosis plots for M1.001 type III signatures (please see *Anticipated Results: Task 4*). For further information about diagnosis plots, please see *Overview of the Chemical Checker data integration pipeline - Interpreting the results: diagnosis plots* and check our Gitlab repository.

### Supplementary code

#### Task 1: Extending a preexisting CC space with additional bioactivity data (B1.002)

```
### DOWNLOADING BIOACTIVITY DATA ###

import os
import wget
import tarfile

# Gathering B1.002 source data
# Modify PATH at will
PATH_TO_InDATA = "../data/"
link = "https://zenodo.org/records/14000635/files/B1.tar.gz?download=1"

# Create path
os.makedirs(PATH_TO_InDATA, exist_ok=True)

# Download
def download_data(PATH_TO_InDATA):
    os.chdir(PATH_TO_InDATA)
    wget.download(link, out=PATH_TO_InDATA )

download_data(PATH_TO_InDATA)

def decompress_data(PATH_TO_FILE, PATH_TO_InDATA):
    os.makedirs(PATH_TO_InDATA, exist_ok=True)
    with tarfile.open(PATH_TO_FILE, "r:gz") as tar:
        tar.extractall(path=PATH_TO_InDATA)

# Modify PATHS at will
PATH_TO_FILE = os.path.join(PATH_TO_InDATA, "B1.tar.gz")
decompress_data(PATH_TO_FILE, PATH_TO_InDATA)
```

```
### FIRST STEPS ###

# Specify the location of the CC config file.
# os.environ['CC_CONFIG'] = '/path/to/your_cc_config.json'  #  e.g. chemicalchecker/setup/cc_config.json
os.environ['CC_CONFIG'] = '/aloy/home/acomajuncosa/cc_config.json'

from chemicalchecker import ChemicalChecker
ChemicalChecker.set_verbosity('DEBUG') # CRITICAL, ERROR, WARN, INFO or DEBUG
from chemicalchecker.core import DataSignature
import numpy as np
import pandas as pd
import json
%matplotlib inline

local_cc_dir = '../local_CC_B1'
PATH_TO_DATA = "/aloy/home/acomajuncosa/CC_DATA/DATA/"  # See Download_Data.ipynb // Procedure step 3
# PATH_TO_DATA = "/aloy/web_checker/package_cc/2021_07/sign_model_links/"
cc_local = ChemicalChecker(local_cc_dir, dbconnect=False, custom_data_path=PATH_TO_DATA)
```

```
### LOADING BIOACTIVITY DATA ###

# Dataset Name
dataset = 'B1.002'

# Input file
inputFile = "../data/B1/repohub_pairs.h5"

# Get old pairs (INCHIKEY-FEATURE)
preprocess_old = "../data/B1/B1.001_preprocess.old.h5"
old_pairs = DataSignature(preprocess_old).get_h5_dataset('pairs')
old_pairs = set([tuple(i) for i in old_pairs])

# Get new pairs (INCHIKEY-FEATURE)
new_pairs = DataSignature(inputFile).get_h5_dataset('pairs')
new_pairs = set([tuple(i) for i in new_pairs])

# Merge pairs
merged_pairs = new_pairs.union(old_pairs)
merged_pairs = np.array(sorted(merged_pairs))

print("NUMBER OF COMPOUNDS: " + str(len(set(merged_pairs[:,0])))) # 6944
print("NUMBER OF MoAs: " + str(len(set(merged_pairs[:,1])))) # 4626
```

```
### TYPE 0 SIGNATURES ###

# Dataset Name
dataset = 'B1.002'

# Instantiation of sign0 data structures for the new space: full and reference
sign0 = cc_local.signature(dataset, 'sign0')

# Cleaning both full and reference datasets. This is crucial!
sign0.clear_all()

# Fit sign0
sign0.fit(pairs=np.array(merged_pairs))

# sign0.shape # (6283, 1314)
```

```
### TYPE I SIGNATURES ###

# Dataset Name
dataset = 'B1.002'

# Instantiation of sign0
sign0 = cc_local.signature(dataset, 'sign0')

# Instantiation of sign1
sign1 = cc_local.signature(dataset, 'sign1')

# Cleaning both full and reference datasets. This is crucial!
sign1.clear_all()

# Fitting sign1
sign1.fit(sign0)
```

```
# sign1.shape # (5826, 438)

# Instantiation of sign1
sign1 = cc_local.signature(dataset, 'sign1')

# Instantiation of neig1
neig1 = cc_local.get_signature("neig1", "full", dataset)  # It will take the reference anyway...

# Cleaning both full and reference. This is crucial!
neig1.clear_all()

# Fitting neig1
neig1.fit(sign1)

# neig1.shape # [5826, 1000]
```

```
### TYPE II SIGNATURES ###
# Dataset Name
dataset = 'B1.002'

# Get sign1
sign1 = cc_local.get_signature('sign1', 'full', dataset)

# Get neig1
neig1 = cc_local.get_signature('neig1', 'full', dataset)  # By default, all vs ref

# Instantiation of sign2
sign2 = cc_local.signature(dataset, 'sign2')

# Cleaning both full and reference datasets. This is crucial!
sign2.clear_all()

# Fit sign2 given sign1 & neig1
sign2.fit(sign1, neig1, oos_predictor=False)
# sign2.shape # (5826, 128)
```

```
### TYPE III SIGNATURES ###

# CHECK OVERLAP

# Dataset Name
dataset = 'B1.002'

# Get CC universe
cc_universe = []
for dat in cc_local.datasets:
   if dat != dataset and dat.endswith('001') and "B1" not in dat:
       cc_universe.extend(cc_local.get_signature('sign2', 'full', dat).keys)
cc_universe = set(cc_universe)

# Get sign2
sign2 = cc_local.signature(dataset, 'sign2')

# Get B1.002 molecules
b1_molecules = set(sign2.keys)
```

```python
print("Number of molecules in the CC universe: " + str(len(cc_universe))) # 1009293
print("Number of molecules in B1.002 sign2: " + str(len(b1_molecules))) # 5826
print("Intersection CC & B1.002: " + str(len(cc_universe.intersection(b1_molecules)))) # 5433

# GET DATA

# Dataset Name
dataset = 'B1.002'

# Instantiation of sign3
sign3 = cc_local.signature(dataset, 'sign3')
sign3.clear_all()

# Create a list of sign2 to feed sign3 -- using the 24 CC spaces (without B1.001) and B1.002
sign2_list = list()

# For each CC space
for ds in cc_local.coordinates:
   ds += '.001'
   if ds == 'B1.001':
       ds = 'B1.002'
   sign2_list.append(cc_local.get_signature('sign2', 'full', ds))

# In total, we now have 25 spaces
print(len(sign2_list))

# Get B1.002 sign1
sign1_self = cc_local.signature(dataset, 'sign1')

# Get B1.002 sign2
sign2_self = cc_local.signature(dataset, 'sign2')

# FIT SIGN3

mapp = None
"""
# Alternatively, you can provide your in-house python dictionary to map InchiKey's to InChI's (see
example below)
mapp = {
'LPXQRXLUHJKZIE-UHFFFAOYSA-N': 'InChI=1S/C4H4N6O/c5-4-6-2-1(3(11)7-4)8-10-9-2/h(H4,5,6,7,8,9,10,11)',
'BZKPWHYZMXOIDC-UHFFFAOYSA-N': 'InChI=1S/C4H6N4O3S2/c1-2(9)6-3-7-8-4(12-
3)13(5,10)11/h1H3,(H2,5,10,11)(H,6,7,9)',
'XZWYZXLIPXDOLR-UHFFFAOYSA-N': 'InChI=1S/C4H11N5/c1-9(2)4(7)8-3(5)6/h1-2H3,(H5,5,6,7,8)'
}
"""

# CAUTION: COMPUTATIONALLY DEMANDING STEP - Consider running it in an HPC cluster
sign3.fit(sign2_list, sign2_self, sign1_self, sign2_universe=None, complete_universe="fast",
sign2_coverage=None, dbconnect=False, mapping_dict=mapp)

# sign3.shape # (1009686, 128)
```

##### Task 2: Rebuilding a preexisting CC space with a different strategy of processing the raw data (D1.002)

```
### DOWNLOADING BIOACTIVITY DATA ###

import os
import wget
import tarfile

# Gathering D1.002 source data
# Modify PATH at will
PATH_TO_InDATA = "../data/"
link = "https://zenodo.org/records/13993758/files/D1.tar.gz?download=1"

# Create path
os.makedirs(PATH_TO_InDATA, exist_ok=True)
# Download
def download_data(PATH_TO_InDATA):
   os.chdir(PATH_TO_InDATA)
   wget.download(link, out=PATH_TO_InDATA )

download_data(PATH_TO_InDATA)
def decompress_data(PATH_TO_FILE, PATH_TO_InDATA):
   os.makedirs(PATH_TO_InDATA, exist_ok=True)
   with tarfile.open(PATH_TO_FILE, "r:gz") as tar:
       tar.extractall(path=PATH_TO_InDATA)

# Modify PATHS at will
PATH_TO_FILE = os.path.join(PATH_TO_InDATA, "D1.tar.gz")
decompress_data(PATH_TO_FILE, PATH_TO_InDATA)
```

```
### FIRST STEPS ###

# Specify the location of the CC config file.
# os.environ['CC_CONFIG'] = '/path/to/your_cc_config.json'  #  e.g. chemicalchecker/setup/cc_config.json
os.environ['CC_CONFIG'] = '/aloy/home/acomajuncosa/cc_config.json'

from chemicalchecker import ChemicalChecker
ChemicalChecker.set_verbosity('DEBUG') # CRITICAL, ERROR, WARN, INFO or DEBUG
from chemicalchecker.core import DataSignature
import numpy as np
import pandas as pd
import json
%matplotlib inline

local_cc_dir = '../local_CC_D1'
PATH_TO_DATA = "/aloy/home/acomajuncosa/CC_DATA/DATA/"  # See Download_Data.ipynb // Procedure step 3
# PATH_TO_DATA = "/aloy/web_checker/package_cc/2021_07/sign_model_links/"
cc_local = ChemicalChecker(local_cc_dir, dbconnect=False, custom_data_path=PATH_TO_DATA)
```

```
### LOADING BIOACTIVITY DATA ###
inputFile = "../data/D1/chdir_active_10.h5"
M = np.array(DataSignature(inputFile))
print(M.shape)  # (11066, 12328)
```

```
### TYPE 0 SIGNATURES ###

# Dataset Name
dataset = 'D1.002'

# Instantiation of sign0 data structures for the new space: full and reference
sign0 = cc_local.signature(dataset, 'sign0')

# Cleaning both full and reference datasets. This is crucial!
sign0.clear_all()

# Fit sign0
sign0.fit(data_file=inputFile, sanitizer_kwargs={"max_features": 13000})

# sign0.shape # (11066, 12328)
```

```
### TYPE I SIGNATURES ###

# Dataset Name
dataset = 'D1.002'

# Instantiation of sign0
sign0 = cc_local.signature(dataset, 'sign0')

# Instantiation of sign1
sign1 = cc_local.signature(dataset, 'sign1')

# Cleaning both full and reference datasets. This is crucial!
sign1.clear_all()

# Fitting sign1
sign1.fit(sign0, scale_kwargs=dict(max_keys=13000), pca_kwargs=dict(max_keys=13000))

# sign1.shape # (11065, 3794)

# Instantiation of sign1
sign1 = cc_local.signature(dataset, 'sign1')

# Instantiation of neig1
neig1 = cc_local.get_signature("neig1", "full", dataset)  # It will take the reference anyway...

# Cleaning both full and reference. This is crucial!
neig1.clear_all()

# Fitting neig1
neig1.fit(sign1)

# neig1.shape # (11065, 1000)
```

```
### TYPE II SIGNATURES ###

# Dataset Name
dataset = 'D1.002'

# Get sign1
sign1 = cc_local.get_signature('sign1', 'full', dataset)
```

```python
# Get neig1
neig1 = cc_local.get_signature('neig1', 'full', dataset)  # By default, all vs ref

# Instantiation of sign2
sign2 = cc_local.signature(dataset, 'sign2')

# Cleaning both full and reference datasets. This is crucial!
sign2.clear_all()

# Fit sign2 given sign1 & neig1
sign2.fit(sign1, neig1, oos_predictor=False)

# sign2.shape # (11065, 128)
```

```python
### TYPE III SIGNATURES ###

# CHECK OVERLAP

# Dataset Name
dataset = 'D1.002'

# Get CC universe
cc_universe = []
for dat in cc_local.datasets:
   if dat != dataset and dat.endswith('001') and "D1" not in dat:
       cc_universe.extend(cc_local.get_signature('sign2', 'full', dat).keys)
cc_universe = set(cc_universe)

# Get sign2
sign2 = cc_local.signature(dataset, 'sign2')

# Get D1.002 molecules
d1_molecules = set(sign2.keys)

print("Number of molecules in the CC universe: " + str(len(cc_universe)))
print("Number of molecules in D1.002 sign2: " + str(len(d1_molecules)))
print("Intersection CC & D1.002: " + str(len(cc_universe.intersection(d1_molecules))))

# GET DATA

# Dataset Name
dataset = 'D1.002'

# Instantiation of sign3
sign3 = cc_local.signature(dataset, 'sign3')
sign3.clear_all()

# Create a list of sign2 to feed sign3 -- using the 24 CC spaces (without D1.001) and D1.002
sign2_list = list()

# For each CC space
for ds in cc_local.coordinates:
   ds += '.001'
   if ds == 'D1.001':
       ds = 'D1.002'
   sign2_list.append(cc_local.get_signature('sign2', 'full', ds))
```

```
# In total, we now have 25 spaces
print(len(sign2_list))

# Get D1.002 sign1
sign1_self = cc_local.signature(dataset, 'sign1')

# Get D1.002 sign2
sign2_self = cc_local.signature(dataset, 'sign2')

# FIT SIGN3

mapp = None
"""
# Alternatively, you can provide your in-house python dictionary to map InchiKey's to InChI's (see
example below)
mapp = {
'LPXQRXLUHJKZIE-UHFFFAOYSA-N': 'InChI=1S/C4H4N6O/c5-4-6-2-1(3(11)7-4)8-10-9-2/h(H4,5,6,7,8,9,10,11)',
'BZKPWHYZMXOIDC-UHFFFAOYSA-N': 'InChI=1S/C4H6N4O3S2/c1-2(9)6-3-7-8-4(12-
3)13(5,10)11/h1H3,(H2,5,10,11)(H,6,7,9)',
'XZWYZXLIPXDOLR-UHFFFAOYSA-N': 'InChI=1S/C4H11N5/c1-9(2)4(7)8-3(5)6/h1-2H3,(H5,5,6,7,8)'
}
"""

# CAUTION: COMPUTATIONALLY DEMANDING STEP - Consider running it in an HPC cluster
sign3.fit(sign2_list, sign2_self, sign1_self, sign2_universe=None, complete_universe="fast",
sign2_coverage=None, dbconnect=False, mapping_dict=mapp)

# sign3.shape # (1011251, 128)
```

#### Task 3: Creating a new CC space from a novel data type that fits the current CC universe (D6.001)

```
### DOWNLOADING BIOACTIVITY DATA ###

import os
import wget
import tarfile

# Modify PATH at will
PATH_TO_InDATA = "../data/"
link = "https://zenodo.org/records/14000624/files/D6.tar.gz?download=1"

# Create path
os.makedirs(PATH_TO_InDATA, exist_ok=True)

# Download
def download_data(PATH_TO_InDATA):
    os.chdir(PATH_TO_InDATA)
    wget.download(link, out=PATH_TO_InDATA )

download_data(PATH_TO_InDATA)

def decompress_data(PATH_TO_FILE, PATH_TO_InDATA):
    os.makedirs(PATH_TO_InDATA, exist_ok=True)
    with tarfile.open(PATH_TO_FILE, "r:gz") as tar:
        tar.extractall(path=PATH_TO_InDATA)

# Modify PATHS at will
PATH_TO_FILE = os.path.join(PATH_TO_InDATA, "D6.tar.gz")
decompress_data(PATH_TO_FILE, PATH_TO_InDATA)
```

```
### FIRST STEPS ###

# Specify the location of the CC config file.
# os.environ['CC_CONFIG'] = '/path/to/your_cc_config.json'  #  e.g. chemicalchecker/setup/cc_config.json
os.environ['CC_CONFIG'] = '/aloy/home/acomajuncosa/cc_config.json'

from chemicalchecker import ChemicalChecker
ChemicalChecker.set_verbosity('DEBUG') # CRITICAL, ERROR, WARN, INFO or DEBUG
import numpy as np
import pandas as pd
import json
%matplotlib inline

local_cc_dir = '../local_CC_D6'
PATH_TO_DATA = "/aloy/home/acomajuncosa/CC_DATA/DATA/"  # See Download_Data.ipynb // Procedure step 3
# PATH_TO_DATA = "/aloy/web_checker/package_cc/2021_07/sign_model_links/"
cc_local = ChemicalChecker(local_cc_dir, dbconnect=False, custom_data_path=PATH_TO_DATA)
```

```
### LOADING BIOACTIVITY DATA ###

# Load the raw binary data
# Rows: compounds
# Columns: Uniprot IDs
```

```
inputFile="../data/D6/D6_processed_LOG2FC.tsv"
df=pd.read_csv(inputFile, sep='\t', index_col=0)
print(df.shape)  # (875, 9960)
```

```
### TYPE 0 SIGNATURES ###

# Dataset Name
dataset = 'D6.001'

# Instantiation of sign0 data structures for the new space: full and reference
sign0 = cc_local.signature(dataset, 'sign0')

# Cleaning both full and reference datasets. This is crucial!
sign0.clear_all()

# Fit sign0
sign0.fit(X=df.values, keys=list(df.index), features=list(df.columns))

# sign0.shape # (873, 9960)
```

```
### TYPE I SIGNATURES ###

# Dataset Name
dataset = 'D6.001'

# Instantiation of sign0
sign0 = cc_local.signature(dataset, 'sign0')

# Instantiation of sign1
sign1 = cc_local.signature(dataset, 'sign1')

# Cleaning both full and reference datasets. This is crucial!
sign1.clear_all()

# Fitting sign1
sign1.fit(sign0)

# sign1.shape # (873, 544)

# Instantiation of sign1
sign1 = cc_local.signature(dataset, 'sign1')

# Instantiation of neig1
neig1 = cc_local.get_signature("neig1", "full", dataset)  # It will take the reference anyway...

# Cleaning both full and reference. This is crucial!
neig1.clear_all()

# Fitting neig1
neig1.fit(sign1)

# neig1.shape # [873, 873]
```

```
### TYPE II SIGNATURES ###

```

```
# Dataset Name
dataset = 'D6.001'

# Get sign1
sign1 = cc_local.get_signature('sign1', 'full', dataset)

# Get neig1
neig1 = cc_local.get_signature('neig1', 'full', dataset)  # By default, all vs ref

# Instantiation of sign2
sign2 = cc_local.signature(dataset, 'sign2')

# Cleaning both full and reference datasets. This is crucial!
sign2.clear_all()

# Fit sign2 given sign1 & neig1
sign2.fit(sign1, neig1, oos_predictor=False)

# sign2.shape # (873, 128)
```

```
### TYPE III SIGNATURES ###

# CHECK OVERLAP

# Dataset Name
dataset = 'D6.001'

# Get CC universe
cc_universe = []
for dat in cc_local.datasets:
    if dat != dataset and dat.endswith('001'):
        cc_universe.extend(cc_local.get_signature('sign2', 'full', dat).keys)
cc_universe = set(cc_universe)

# Get sign2
sign2 = cc_local.signature(dataset, 'sign2')

# Get D6 molecules
d6_molecules = set(sign2.keys)

print("Number of molecules in the CC universe: " + str(len(cc_universe))) # 1009293
print("Number of molecules in D6 sign2: " + str(len(d6_molecules))) # 873
print("Intersection CC & D6: " + str(len(cc_universe.intersection(d6_molecules)))) # 803

# GET DATA

# Instantiation of sign3
sign3 = cc_local.signature(dataset, 'sign3')
sign3.clear_all()

# Create a list of sign2 to feed sign3 -- using the 25 CC spaces & D6
sign2_list = list()

# For each CC space
for ds in cc_local.coordinates:
    ds += '.001'
    sign2_list.append(cc_local.get_signature('sign2', 'full', ds))
```

```
# Append the new D6 space
sign2_list.append(cc_local.get_signature('sign2','full', dataset))

# In total, we now have 26 spaces
print(len(sign2_list)) # 26

# Get D6 sign1
sign1_self = cc_local.signature(dataset, 'sign1')

# Get D6 sign2
sign2_self = cc_local.signature(dataset, 'sign2')

# FIT SIGN3

mapp = None
"""
# Alternatively, you can provide your in-house python dictionary to map InchiKey's to InChI's (see
example below)
mapp = {
'LPXQRXLUHJKZIE-UHFFFAOYSA-N': 'InChI=1S/C4H4N6O/c5-4-6-2-1(3(11)7-4)8-10-9-2/h(H4,5,6,7,8,9,10,11)',
'BZKPWHYZMXOIDC-UHFFFAOYSA-N': 'InChI=1S/C4H6N4O3S2/c1-2(9)6-3-7-8-4(12-
3)13(5,10)11/h1H3,(H2,5,10,11)(H,6,7,9)',
'XZWYZXLIPXDOLR-UHFFFAOYSA-N': 'InChI=1S/C4H11N5/c1-9(2)4(7)8-3(5)6/h1-2H3,(H5,5,6,7,8)'
}
"""

# CAUTION: COMPUTATIONALLY DEMANDING STEP - Consider running it in an HPC cluster
sign3.fit(sign2_list, sign2_self, sign1_self, sign2_universe=None, complete_universe="fast",
sign2_coverage=None, dbconnect=False, mapping_dict=mapp)

# sign3.shape # (1009363, 128)
```

##### Task 4: Creating a new CC space from a novel data type that does not fit the current CC universe (M1.001)

```
### DOWNLOADING BIOACTIVITY DATA ###

import os
import wget
import tarfile

# Gathering M1 source data
# Modify PATH at will
PATH_TO_InDATA = "../data/"
link = "https://zenodo.org/records/14000612/files/M1.tar.gz?download=1"

# Create path
os.makedirs(PATH_TO_InDATA, exist_ok=True)

# Download
def download_data(PATH_TO_InDATA):
    os.chdir(PATH_TO_InDATA)
    wget.download(link, out=PATH_TO_InDATA )

download_data(PATH_TO_InDATA)

def decompress_data(PATH_TO_FILE, PATH_TO_InDATA):
    os.makedirs(PATH_TO_InDATA, exist_ok=True)
    with tarfile.open(PATH_TO_FILE, "r:gz") as tar:
        tar.extractall(path=PATH_TO_InDATA)

# Modify PATHS at will
PATH_TO_FILE = os.path.join(PATH_TO_InDATA, "M1.tar.gz")
decompress_data(PATH_TO_FILE, PATH_TO_InDATA)
```

```
### FIRST STEPS ###

# Specify the location of the CC config file.
# os.environ['CC_CONFIG'] = '/path/to/your_cc_config.json'  #  e.g. chemicalchecker/setup/cc_config.json
os.environ['CC_CONFIG'] = '/aloy/home/acomajuncosa/cc_config.json'

from chemicalchecker import ChemicalChecker
ChemicalChecker.set_verbosity('DEBUG') # CRITICAL, ERROR, WARN, INFO or DEBUG
import numpy as np
import pandas as pd
import json
%matplotlib inline

local_cc_dir = '../local_CC_M1'
PATH_TO_DATA = "/aloy/home/acomajuncosa/CC_DATA/DATA/"  # See Download_Data.ipynb // Procedure step 3
# PATH_TO_DATA = "/aloy/web_checker/package_cc/2021_07/sign_model_links/"
cc_local = ChemicalChecker(local_cc_dir, dbconnect=False, custom_data_path=PATH_TO_DATA)
```

```
### LOADING BIOACTIVITY DATA ###

inputFile="../data/M1/microbiota_raw.csv"
df=pd.read_csv(inputFile,index_col=0)
```

```
print(df.shape)  # (1155, 40)
```

```
### TYPE 0 SIGNATURES ###

# Dataset Name
dataset = 'M1.001'

# Instantiation of sign0 data structures for the new space: full and reference
sign0 = cc_local.signature(dataset, 'sign0')

# Cleaning both full and reference datasets. This is crucial!
sign0.clear_all()

# Fit sign0
sign0.fit(X=df.values, keys=list(df.index), features=list(df.columns))

# sign0.shape # (310, 40)
```

```
### TYPE I SIGNATURES ###

# Dataset Name
dataset = 'M1.001'

# Instantiation of sign0
sign0 = cc_local.signature(dataset, 'sign0')

# Instantiation of sign1
sign1 = cc_local.signature(dataset, 'sign1')

# Cleaning both full and reference datasets. This is crucial!
sign1.clear_all()

# Fitting sign1
sign1.fit(sign0)

# sign1.shape # (200, 13)

# Instantiation of sign1
sign1 = cc_local.signature(dataset, 'sign1')

# Instantiation of neig1
neig1 = cc_local.get_signature("neig1", "full", dataset)

# Cleaning both full and reference. This is crucial!
neig1.clear_all()

# Fitting neig1
neig1.fit(sign1)

# neig1.shape # [200, 177]
```

```
### TYPE II SIGNATURES ###

# Dataset Name
dataset = 'M1.001'
```

```
# Get sign1
sign1 = cc_local.get_signature('sign1', 'full', dataset)

# Get neig1
neig1 = cc_local.get_signature('neig1', 'full', dataset)  # By default, all vs ref

# Instantiation of sign2
sign2 = cc_local.signature(dataset, 'sign2')

# Cleaning both full and reference datasets. This is crucial!
sign2.clear_all()

# Fit sign2 given sign1 & neig1
sign2.fit(sign1, neig1, oos_predictor=False)

# sign2.shape # (200, 128)
```

```
### TYPE III SIGNATURES ###

# CHECK OVERLAP

# Dataset Name
dataset = 'M1.001'

# Get CC universe
cc_universe = []
for dat in cc_local.datasets:
    if dat != dataset and dat.endswith('001'):
        cc_universe.extend(cc_local.get_signature('sign2', 'full', dat).keys)
cc_universe = set(cc_universe)

# Get sign2
sign2 = cc_local.signature(dataset, 'sign2')

# Get M1 molecules
m1_molecules = set(sign2.keys)

print("Number of molecules in the CC universe: " + str(len(cc_universe))) # 1009293
print("Number of molecules in M1 sign2: " + str(len(m1_molecules))) # 200
print("Intersection CC & M1: " + str(len(cc_universe.intersection(m1_molecules)))) # 188

# GET DATA

# Instantiation of sign3
sign3 = cc_local.signature(dataset, 'sign3')
sign3.clear_all()

# Create a list of sign2 to feed sign3 -- using the 25 CC spaces & M1
sign2_list = list()

# For each CC space
for ds in cc_local.coordinates:
    ds += '.001'
    sign2_list.append(cc_local.get_signature('sign2', 'full', ds))

# Append the new M1 space
sign2_list.append(cc_local.get_signature('sign2','full', dataset))
```

```
# In total, we now have 26 spaces
print(len(sign2_list)) # 26

# Get M1 sign1
sign1_self = cc_local.signature(dataset, 'sign1')

# Get M1 sign2
sign2_self = cc_local.signature(dataset, 'sign2')

# FIT SIGN3

mapp = None
"""
# Alternatively, you can provide your in-house python dictionary to map InchiKey's to InChI's (see
example below)
mapp = {
'LPXQRXLUHJKZIE-UHFFFAOYSA-N': 'InChI=1S/C4H4N6O/c5-4-6-2-1(3(11)7-4)8-10-9-2/h(H4,5,6,7,8,9,10,11)',
'BZKPWHYZMXOIDC-UHFFFAOYSA-N': 'InChI=1S/C4H6N4O3S2/c1-2(9)6-3-7-8-4(12-
3)13(5,10)11/h1H3,(H2,5,10,11)(H,6,7,9)',
'XZWYZXLIPXDOLR-UHFFFAOYSA-N': 'InChI=1S/C4H11N5/c1-9(2)4(7)8-3(5)6/h1-2H3,(H5,5,6,7,8)'
}
"""

# CAUTION: COMPUTATIONALLY DEMANDING STEP - Consider running it in an HPC cluster
sign3.fit(sign2_list, sign2_self, sign1_self, sign2_universe=None, complete_universe="fast",
sign2_coverage=None, dbconnect=False, mapping_dict=mapp)

# sign3.shape # (1009305, 128)
```
